# Single-cell transcriptome imaging reveals conserved and virulence-linked phenotypic states in bacteria

**DOI:** 10.64898/2026.02.01.703127

**Authors:** Zohar Persky, Camilla Ciolli Mattioli, Knan Ztion Garcia, Ofir Azran, Roi Avraham, Daniel Dar

## Abstract

Single-cell phenotypic diversity underlies fundamental bacterial behaviors. Resolving this heterogeneity at scale while maintaining sensitive detection remains a central challenge. Here, we address this gap by combining the strengths of single-molecule fluorescence *in situ* hybridization (smFISH) and transcriptome-wide sequencing. We present par^2^FISH, a framework for discovering transcriptionally defined subpopulations across conditions by multiplexing tens of thousands of smFISH reactions. Through comparative single-cell transcriptomics in *Escherichia coli* and *Salmonella*, we uncover conserved and species-specific patterns of heterogeneity, including mutually exclusive states, coordinated metabolic specialization, and differential activation of virulence programs. We further obtain complete transcriptomes for selected subpopulations using marker-based cell sorting and RNA-sequencing (FACS-RNA-seq). This integrated strategy paves the way for exposing bacterial phenotypic landscapes and the environmental, physiological, and evolutionary factors that shape them.

## Background

Individual bacteria can exhibit distinct phenotypes despite sharing the same genotype and environment (*1–4*). Such non-genetic heterogeneity arises from stochastic fluctuations in gene expression and regulatory circuits that amplify or stabilize these fluctuations, generating functionally distinct subpopulations (*1*, *5*, *6*). Phenotypic diversity is thought to enhance robustness to environmental perturbations, including antibiotic exposure (*7*, *8*), and to enable cooperative division of labor underlying ecological success and virulence (*1*, *3*, *9–11*). Despite its apparent importance, the identities of discrete cellular states, their functional roles, and the regulatory principles governing heterogeneity remain incompletely understood, even in model species. Addressing these gaps requires methods that can systematically and sensitively quantify gene expression at the single-cell level.

The single-bacterium transcriptomics toolbox has expanded rapidly over the past several years (*12–20*). Nevertheless, substantial technical challenges continue to limit progress. Chief among these is the extremely low abundance of bacterial mRNAs, typically fewer than one transcript per gene per cell (*21*, *22*), their lack of polyadenylation, and their encapsulation within rigid cell walls that impede efficient extraction (22). In addition, total transcriptome size can decrease markedly as a function of growth phase and doubling rate (*23*, *24*), further compounding sensitivity constraints. Recent advances have begun to address these barriers through approaches that can be broadly grouped into two classes: RNA sequencing based methods (*12–18*) and imaging based platforms (*19*, *20*).

RNA sequencing offers transcriptome-wide coverage but currently achieves relatively low capture efficiencies and exhibits sampling biases toward highly expressed genes (*12–18*). In addition, these destructive techniques do not preserve cellular or spatial information. In contrast, imaging-based approaches such as parallel and sequential FISH (par-seqFISH) and multiplexed error-robust FISH (MERFISH) preserve physical context but measure a targeted set of genes (*19*, *20*). In par-seqFISH, only 100 to 150 genes are measured within the same cell, yet transcript detection efficiency is maximized through the use of smFISH, the most sensitive approach currently available, and dozens of conditions can be screened simultaneously via combinatorial cell labeling (*19*, *23*). Transcript barcoding approaches in physically expanded cells, such as MERFISH, increase multiplexing capacity by approximately an order of magnitude but incur substantial losses in detection efficiency and in the number of analyzed cells as a function of the number of measured genes (*20*). Since no single modality can simultaneously maximize gene coverage and detection sensitivity, we sought to develop an integrated platform that combines the complementary strengths of imaging and transcriptome sequencing.

Here, we present a multistep framework grounded in the observation that bacterial subpopulations can be defined by marker genes exhibiting exceptionally high cell-to-cell variability, analogous to cell-type markers in eukaryotic systems. To maximize detection sensitivity, we developed par^2^FISH, which multiplexes tens of thousands of smFISH reactions to screen thousands of genes across diverse conditions and identify subpopulation markers. We then introduce FISH-Links, a complementary approach that reconstructs subpopulation-level transcriptional programs from separate smFISH measurements and can be applied to characterize dozens of subpopulations in parallel. We further developed a low-input, cell sorting based RNA sequencing approach to obtain comprehensive transcriptomes for subpopulations identified by par^2^FISH.

We applied this strategy to analyze over 4,300 genes across dozens of conditions in two key bacterial species, the model organism *E. coli* and the gastroenteric pathogen *Salmonella enterica* serovar Typhimurium (*S. Typhimurium*). This analysis resolves the phenotypic structure of these bacteria at unprecedented resolution, capturing previously described subpopulations while revealing diverse new states. Through comparative single-cell transcriptomics, we uncover conserved principles and species-specific features of bacterial heterogeneity, including mutually exclusive states and coordinated metabolic specialization. In *S. Typhimurium*, we provide new insights into how cell-to-cell variability in virulence factor expression is regulated and coupled to metabolic reprogramming through environmental sensing. Together, these results establish a general approach that maximizes discovery potential and depth of characterization for dissecting bacterial phenotypic landscapes and their regulatory and evolutionary foundations.

## Results

### Scaling par-seqFISH to the transcriptome level

Par-seqFISH uses high-sensitivity smFISH to profile single-cell gene expression across multiple conditions but is limited to 100-150 genes per experiment (*19*, *23*). To overcome this limitation, we developed par^2^FISH (for parallel par-seqFISH), a multiplexed imaging strategy that scales smFISH to transcriptome-level gene coverage. In this approach, cells collected from multiple conditions are fixed and hybridized with barcoding probes targeting the 16S rRNA to encode their condition of origin and then pooled into a single mixture. The pooled cells are subdivided into multiple hybridization reactions, each containing a distinct probe library targeting a different subset of genes, such that each reaction constitutes an independent par-seqFISH experiment (Fig. 1A). A second barcode is introduced at this stage to record probe library identity (Fig. 1A). As a result, each cell carries two barcodes: one specifying its original growth condition and another identifying the gene panel with which it was probed. Following hybridization, cells are pooled again and imaged to decode both barcodes and quantify gene expression across all panels (Fig. 1B).

**Fig. 1.**
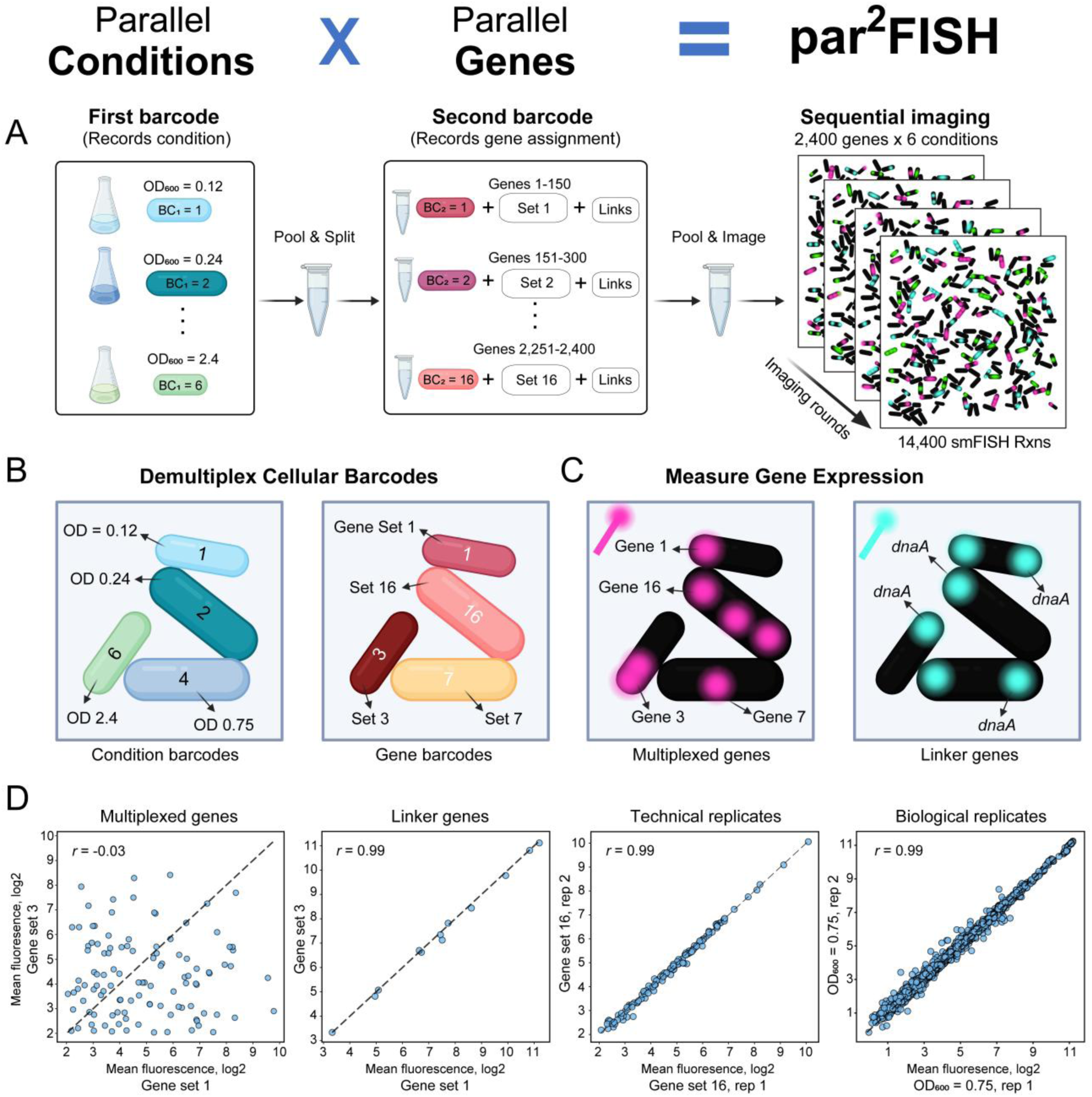
Massively multiplexed smFISH using par^2^FISH. (**A**) Experimental workflow of par^2^FISH. Bacterial cultures are sampled at different growth stages and barcoded with 16S rRNA binding probes that record the growth condition (first barcode; BC_1_). Samples are pooled and split into groups, each hybridized with a distinct gene probe library and a second 16S rRNA barcode (BC_2_) that records the gene assignment. Imaging is performed using automated microscopy in sequential rounds (2,400 genes x 6 conditions, totaling 14,400 smFISH reactions in a single imaging session). (**B**) Dual barcoding encodes both the growth condition and gene library assignment, enabling pooled imaging and subsequent demultiplexing of single cells. (**C**) Each library contains a set of 150 unique genes and a shared set of linker genes. After demultiplexing, specific genes are assigned to each fluorescent readout and cell. For linker genes, each readout measures the same gene across all cells. (**D**) Pairwise comparisons showing reproducibility across (1) multiplexed genes and (2) linker genes between different gene sets, (3) technical replicates (same gene set, different barcodes), and (4) biological replicates (same conditions, two independent samples and barcodes, showing all measured genes). The Pearson correlation values are shown in the figures. Cartoons were created with BioRender.

To validate this method, we designed 16 unique probe libraries targeting a combined total of 2,400 *E. coli* genes, representing about 64% of all known transcriptional units in this organism (Table S1). Additionally, each probe library included a set of shared reference genes measured in every cell that provide internal controls for probe synthesis and hybridization efficiency (Fig. 1C). We term these genes “linkers” for their ability to connect separate experiments (as described later). The chosen linker genes included several known subpopulation marker genes, including *fimA* (*15*, *25*), *gadB* (*15*, *18*, *26*), and *fliC* (*15*, *27*). *E. coli* cultures were grown in M9+0.2% glucose medium, and samples were collected and fixed at five optical densities ranging from OD_600_ of 0.12 to 2.4 (Fig. 1A-B). Samples were barcoded in two steps, as described above. A negative control, as well as technical and biological replicates were included. Under this design, each fluorescent probe reads 96 different measurements (16 genes x 6 conditions) per imaging round, generating a total of 14,400 potential smFISH data points in this par^2^FISH experiment (Fig. 1A).

We demultiplexed >282,000 cells with estimated library misclassification rate of 0.07% and an average of 2,615 cells per library-condition combination. Linker gene signals were highly correlated across libraries (*r* = 0.99, *p* < 10^-10^; Fig. 1D), whereas multiplexed genes showed no meaningful correlation (average *r* = 0.04; Fig. 1D), supporting high signal efficiency and specificity. Replicate libraries encoded by different barcodes, as well as full biological replicates, exhibited nearly identical average expression profiles, underscoring high technical reproducibility (*r* = 0.99, *p* < 10^-10^; Fig. 1D). Furthermore, comparing the average smFISH signals to previously published RNA-sequencing data collected under similar conditions (*28*), showed strong agreement (*r* = 0.79; *p* < 10^-10^; fig. S1), on par with previous comparisons of multiplexed FISH and RNA-seq data (*19*, *20*, *29*). Together, these results establish par^2^FISH as a robust and scalable method for transcriptome-level single-cell expression analysis.

### Mapping the single-cell phenotypic landscape of *E. coli*

The *E. coli* phenotypic landscape has been extensively characterized through decades of focused studies (*10*, *25–27*), large-scale fluorescent reporter screens (*30*, *31*), and more recently, using scRNA-seq (*15*, *18*). This vast knowledgebase provides a strong reference for evaluating par^2^FISH as a tool to uncover functional subpopulations systematically. To identify subpopulation marker genes, we integrated two metrics of expression variability: the coefficient of variation (CV) and the Tails Expression Ratio (TER) (*23*). Since variability depends on mean expression, we plotted log-transformed CV and TER values against their respective means, fitted the relationships, and quantified residual variability as the deviation from the fit (Fig. 2A–B). Residuals from both metrics were then standardized (z-scored) and averaged to produce a unified Expression Variability Index (EVI), which was highly correlated between replicates (*r* = 0.95; *p* < 10^-10^; fig. S2). Notably, the median EVI score was similar across optical densities, indicating a lack of global changes in expression heterogeneity in these conditions (Fig. 2A). Gene-condition datapoints were ranked by EVI, and the top 1% were classified as hypervariable. This analysis identified 41 genes with exceptional cell-to-cell variability, about 70% of which were previously described (Table S2). These included genes involved in stress responses, physiology, and auxiliary functions, several of which we independently validated using promoter reporters, and are described below (Fig. 2C–E; Table S2; fig. S3).

**Fig. 2.**
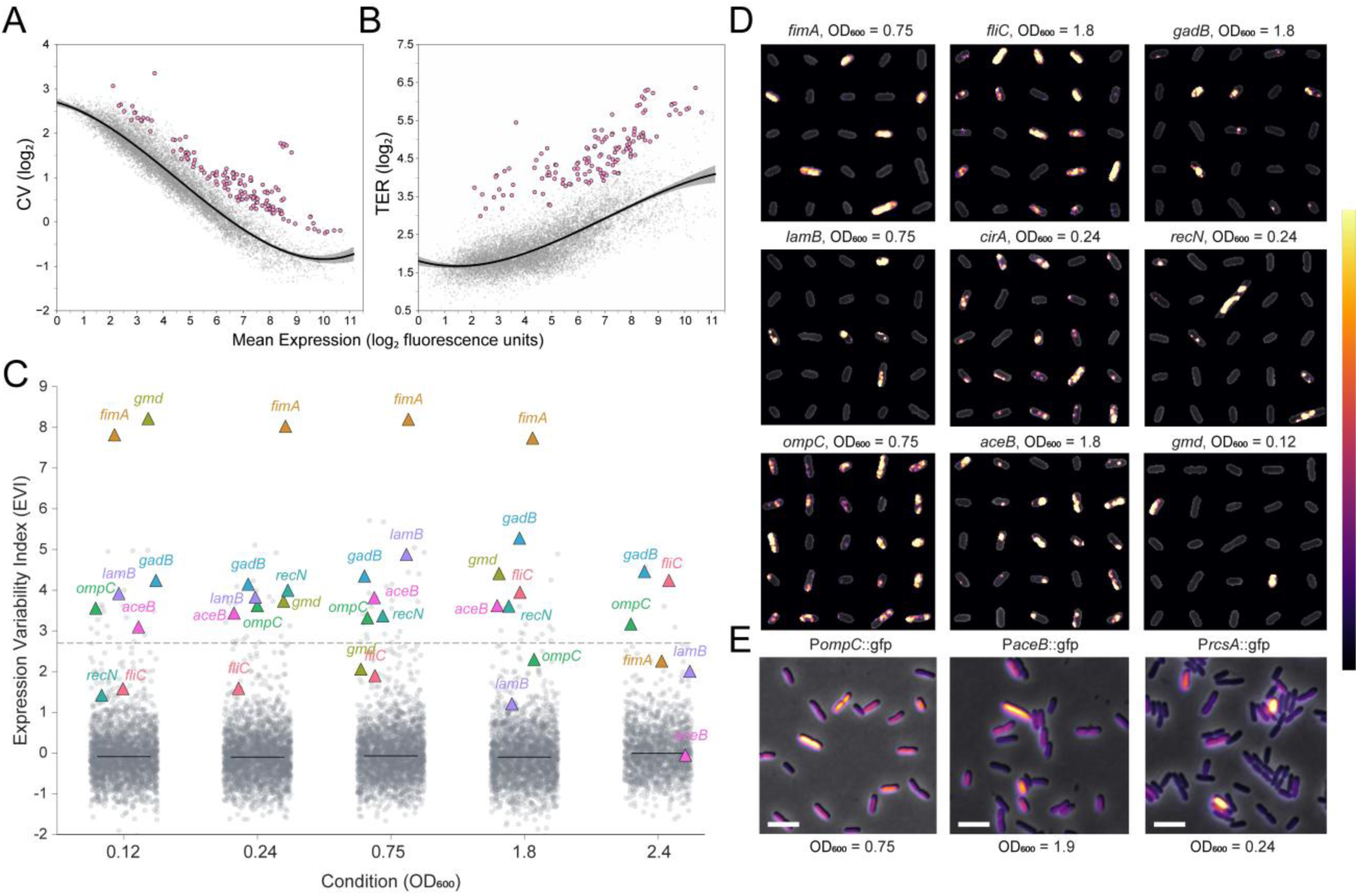
Dissecting the phenotypic landscape of *E. coli* with par^2^FISH. (**A-B**) Coefficient of variation (CV) and Tails Expression Ratio (TER) values plotted against mean expression for each gene–condition datapoint with expression in a sufficient number of cells (n = 13,006). Individual datapoints are shown in gray, and those passing the heterogeneity threshold based on integrated CV and TER metrics (EVI score) are highlighted in pink. Mean expression values are reported in arbitrary fluorescence units. The fitted trend and 95% confidence intervals are shown as a black line and gray shading, respectively. (**C**) EVI distributions per condition (EVI calculated with datapoints from all conditions). The threshold EVI demarking the top 1% of variable data points is shown as a dashed line (EVI = 2.7). The median EVI of each condition is shown as a horizontal black line. All gene-condition data points are shown as gray points with several genes of interest highlighted as colored triangles. (**D**) Raw par^2^FISH data shown over randomly selected cells for several hypervariable genes. (**E**) Independent validation using promoter reporters for three genes (*ompC*, *aceB*, and *rcsA*, the activator of *gmd*); additional reporters shown in fig. S3. Scale bar, 5 µm. Fluorescence intensity (D-E) is displayed with the inferno color map, as shown in the reference bar within the figure.

Bacteria can remodel their surfaces with external structures such as fimbriae, flagella, and capsules to promote adhesion, motility, and immune evasion; functions previously linked to regulated phenotypic heterogeneity. Indeed, the most hypervariable gene in our dataset was *fimA*, encoding the type 1 fimbriae subunit (Fig. 2C-D). This gene is controlled by a genetic phase variation mechanism, in which a promoter inversion generates subpopulations with strong and bistable “on”/“off” expression states (*25*). Similarly, expression of the flagellin, *fliC*, was restricted to subpopulations, especially at higher densities (Fig. 2C-D), consistent with recent reports (*15*). *E. coli* can produce the protective colanic acid capsule (M antigen). Strikingly, two biosynthetic genes from this pathway, *gmd* and *wcaI* (of the same operon), ranked among the top variable genes, showing high expression in a rare subpopulation during early exponential growth (Fig. 2C-D; Table S2). Notably, *rcsA*, the transcriptional activator of these genes, also ranked highly in a previous largescale promoter reporter screen (*30*), and was independently validated here, supporting the formation of encapsulated and protected minority (Fig. 2E).

Phenotypic heterogeneity has also been extensively linked to stress responses and bet-hedging scenarios (*1*, *3*, *4*, *32*). For example, during glucose fermentation, *E. coli* acidifies its environment and activates glutamate-dependent acid resistance genes in a subset of cells, in anticipation of further acidification (*18*, *26*). Indeed, we identify *gadB* and related acid-stress genes (*gadA*, *gadC*, and *hdeA*) among the top hypervariable loci (Fig. 2C-D; Table S2). Similarly, in agreement with reports of pulsatile activation of the DNA-damage SOS response (*33*), we identify subpopulation-specific expression of *recN* (Fig. 2C-D). Stress adaptation can also involve tuning outer membrane permeability. Strikingly, the outer membrane porin gene *ompC* consistently ranked among the top variable genes across all conditions (Fig. 2C-E). In *Salmonella*, *ompC* expression predicts differential antibiotic resistance (*34*), suggesting a similar role in *E. coli*.

Heterogeneity in nutrient utilization strategies can allow for asynchronous resource consumption and enhance survival in fluctuating environments (*32*, *35*, *36*). During aerobic growth on glucose, *E. coli* performs overflow metabolism, excreting acetate that can later be consumed via the glyoxylate shunt as glucose is depleted (*37*). We find that *aceB*, encoding malate synthase, the key glyoxylate shunt enzyme, shows subpopulation-specific expression (Fig. 2C-E), suggesting co-existing groups may adopt distinct carbon source preferences within the same well-mixed culture. In addition to acetate metabolism, we observed hypervariable expression of the maltose/maltodextrin import genes (*malE* and *lamB*), as well as *srlD*, a gene involved in sorbitol utilization (Fig. 2C-E; Table S2). Given our defined growth medium, these findings point toward specialization in carbon source consumption strategies, even in the absence of specific substrates.

### Reconstructing transcriptional programs from separate smFISH reactions

Phenotypic states can be defined by the coordinated regulation of multiple genes, and in such cases cannot be fully characterized by detection of individual marker genes alone. We therefore developed FISH-Links, a second-stage approach that reconstructs subpopulation-level transcriptional signatures via linker genes measured in follow-up par^2^FISH experiments. In this framework, probes targeting subpopulation markers (e.g., *fimA*) are included in every probe library during hybridization and are therefore measured in all cells (Fig. 1A). Cells are partitioned into Linker^High^ and Linker^Low^ subpopulations based on marker levels, and differential expression is computed for each gene within these groups. Although individual genes are generally measured in different cells, their relative enrichment or depletion across linker-defined subpopulations can be quantified and integrated to reconstruct multigene transcriptional states (Fig. 3A).

**Fig. 3.**
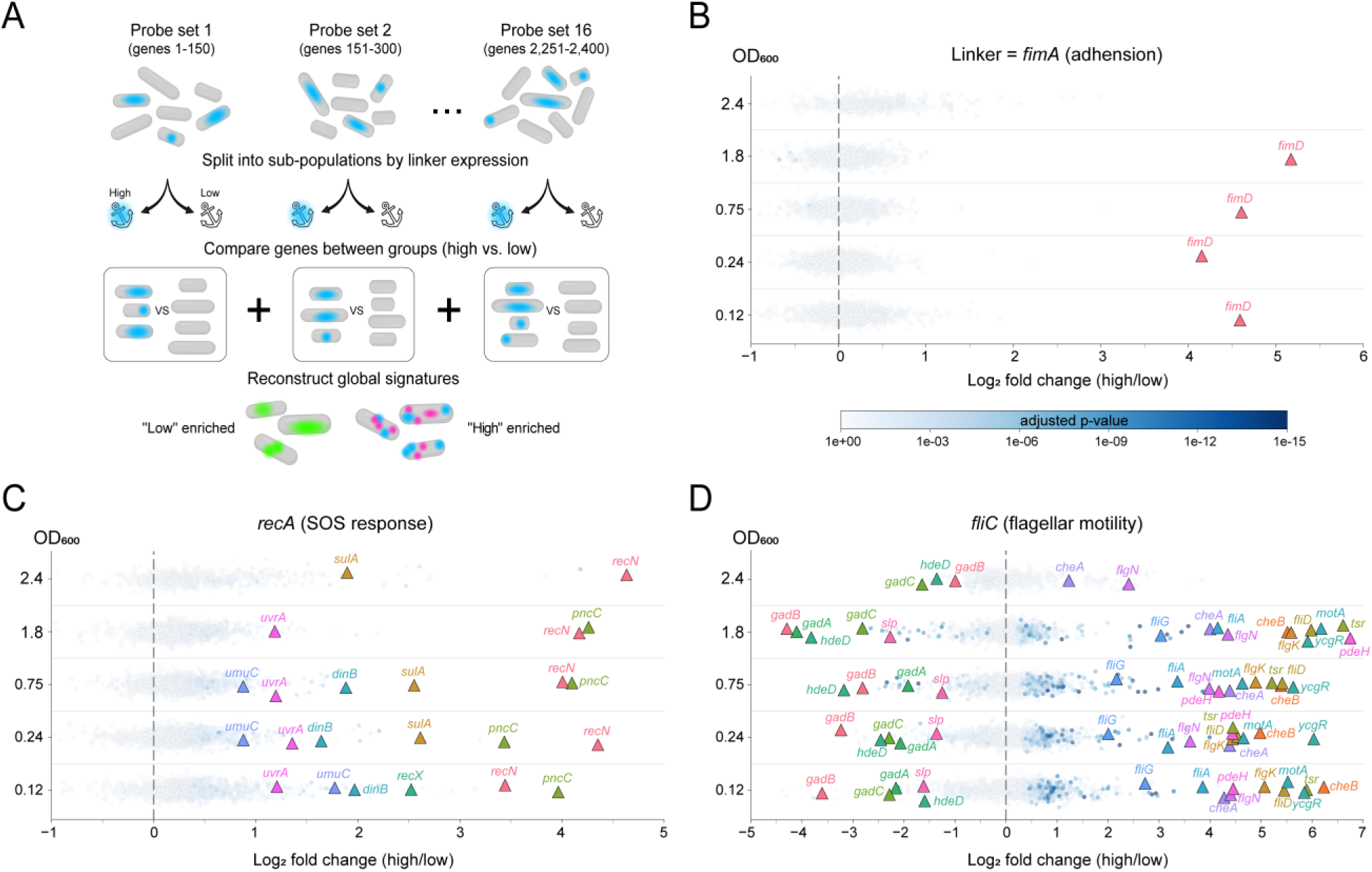
FISH-Links subpopulation transcriptome reconstruction. (**A**) Overview of the FISH-Links approach. Multiplexed gene sets are measured in different cells (in the same par^2^FISH experiment), while linker genes are measured in all cells. Linker expression (blue) is used to split cells *in silico* into high- and low-expressing groups. For each gene, differential expression analysis is performed between the Linker^High^ and Linker^Low^ subpopulations, and the results are integrated to identify genes enriched in each subpopulation. (**B**) FISH-Links analysis using a *fimA* linker. The x-axis shows the log_2_ fold change between *fimA*^High^ and *fimA*^Low^ cells. Genes expressed in a sufficient number of cells in each group are shown as points and are colored according to the p-value (adjusted for multiple hypothesis testing). For visual clarity, p-value color is capped at 10⁻¹⁵. Differentially expressed genes of interest are highlighted as colored triangles. Results from different growth conditions (OD_600_ values) are stacked along the y-axis. (**C–D**) FISH-Links analyses using the SOS master regulator *recA* (**C**) and the flagellin *fliC* (**D**) as linkers. Points are colored as in (B). Cartoons were created with BioRender.

Our initial experiment included several hypervariable linker genes as internal controls, providing a means to evaluate the FISH-Links strategy within the same dataset (Fig. 1). As an initial test case, we compared *fimA*^High^ and *fimA*^Low^ subpopulations, defined as the top and bottom 5% expressing cells (Fig. 3A). Out of thousands of analyzed genes, only *fimD* was highly up-regulated, in agreement with the locus-specific control of the phase variation mechanism (Fig. 3B; Table S3). In a separate example, our screen identified *recN* as a marker for DNA damage in a subset of cells. Indeed, using the SOS master regulator gene, *recA,* as a linker revealed coordinated upregulation of the key SOS genes (*recN*, *pncC*, *sulA*, etc.) within the same subpopulation (Fig. 3C; Table S3). Next, we sought to understand why *fliC*, encoding the major flagellin, was only expressed in a subset of cells. Within *fliC*^High^ cells we uncovered induction of diverse flagellar and chemotaxis genes, including their regulators, supporting differential motility (Fig. 3D). Strikingly, *fliC*^High^ cells strongly downregulated the acid-stress response genes (e.g., *gadB*) up to 16-fold with differences between predicted motile and non-motile subpopulations peaking at OD_600_=1.8 (Fig. 3D; Table S3). We validated this mutual exclusivity using non-multiplexed smFISH and found that it also emerges in early stationary phase (ESP) LB cultures (fig. S4). These coexisting and mutually exclusive states are consistent with a recently described regulatory link (*38*) and support a bet-hedging model in which cells either commit preemptively to the future stress of acidification or maintain motility to explore new environments.

### Comparative analysis reveals conserved and species-specific heterogeneity

While phenotypic heterogeneity has been extensively characterized within individual bacterial species, its conservation across organisms has not been systematically examined at single-cell resolution. Identifying conserved heterogeneity patterns may reveal fundamental cellular behaviors and guide prioritization of subpopulation characterization. Par^2^FISH provides a quantitative framework for such comparisons. We therefore extended our analysis to *S*. *Typhimurium*, which shares an ecological niche, as well as many of its genes with *E. coli*, but has diverged into a distinct pathogenic lifestyle. To make the most direct comparison, we grew *S. Typhimurium* in identical culturing conditions and collected samples at comparable densities to our *E. coli* dataset. We targeted >1,920 genes (Table S4) using par^2^FISH and demultiplexed >208,000 cells. Quantifying single-cell expression, we identified 76 hypervariable genes, using a slightly relaxed threshold as described below (fig. S5; Table S5).

To compare expression variability between *E. coli* and *S. Typhimurium*, we analyzed the maximum EVI scores of 826 orthologous genes measured with sufficient coverage under matched conditions (Fig. 4A). Variability patterns were significantly correlated (*r* = 0.44, *p* < 10^-10^). Binary classification of genes as hypervariable or not revealed strong concordance between the two species: genes hypervariable in *E. coli* were 28.5 times more likely to be hypervariable in *S. Typhimurium* than expected by chance (Fisher’s exact test, *p* = 3.7 x 10^-8^). A modest asymmetry was observed (McNemar’s *p* = 0.04), with hypervariability occurring more frequently in *S. Typhimurium*. This bias was largely attributable to flagellar and chemotaxis genes: in *S. Typhimurium*, a striking 26% of hypervariable genes (20/76) mapped to these functions (Table S5), with elevated heterogeneity evident across most growth conditions (hence our more relaxed EVI threshold in this organism). In contrast, in *E. coli* only *fliC* reached the hypervariability threshold (Table S2), and only in late exponential phase. This difference reflects the stronger positive-feedback bistability circuit in *S. Typhimurium* (*ydiV*-*fliZ* regulatory loop), which is weakly configured in *E. coli*, producing dynamic pulses instead of stable states (*27*, *39*).

**Fig. 4.**
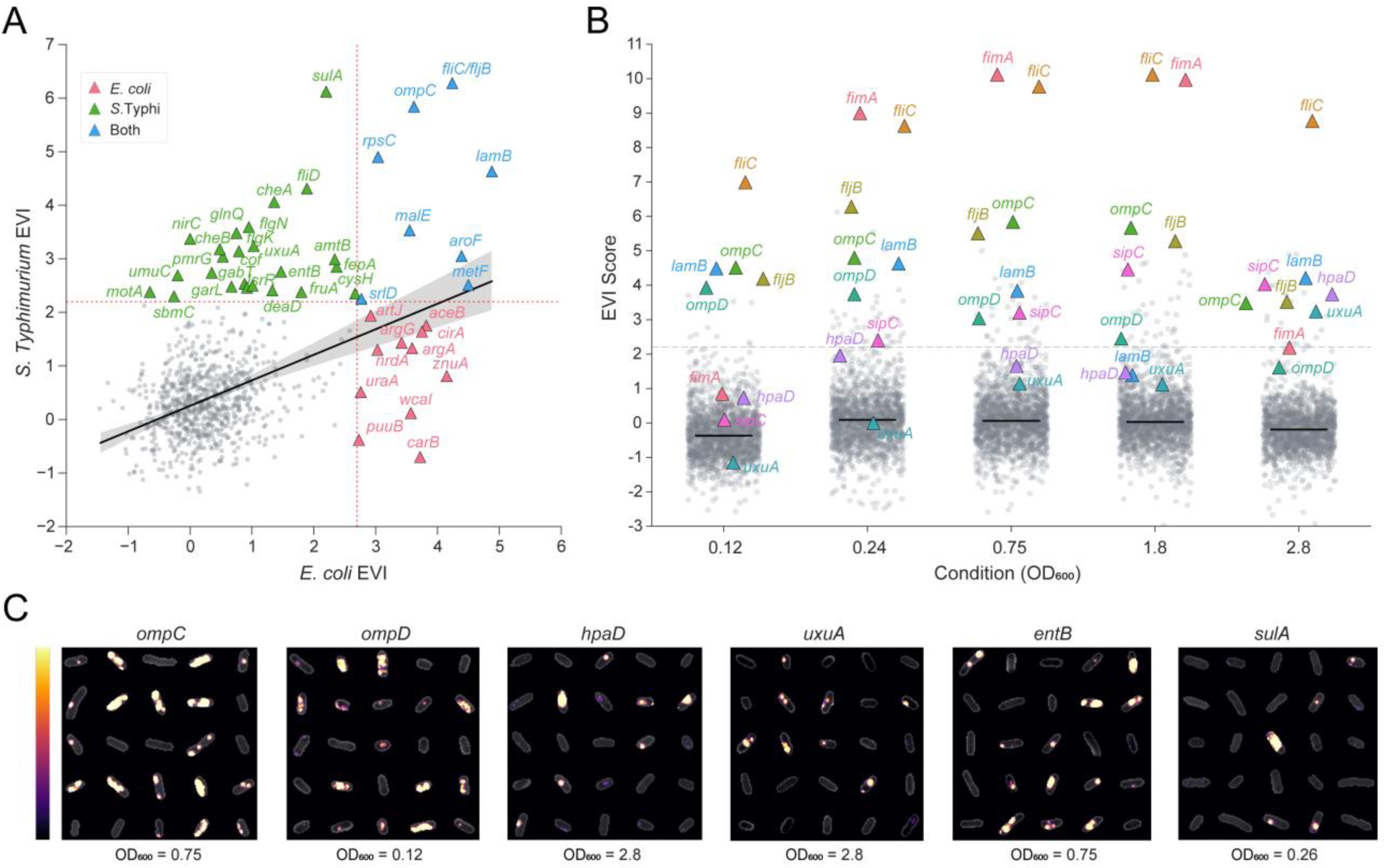
Comparative single-cell transcriptomics of *E. coli* and *S. Typhimurium*. Comparative analysis of *E. coli* and *S. Typhimurium* grown under identical conditions and population densities. (**A**) Scatter plot showing maximal EVI scores for 826 orthologous genes. Species-specific EVI thresholds for hypervariability are indicated by vertical and horizontal red dotted lines. Gray points represent genes below the threshold in both species, and colored triangles denote hypervariable genes in one or both species (see inset color legend). A fitted linear regression is shown in black with the 95% confidence interval in gray (Pearson’s *r* = 0.44, *p* < 10^-10^). (**B**) Distribution of *S. Typhimurium* EVI values per condition as indicated in the x-axis (EVI calculated using datapoints from all conditions). A dashed line marks the EVI hypervariability threshold (EVI = 2.21). Median EVI per condition is shown as a horizontal black line; all gene–condition datapoints are shown in gray, with selected genes highlighted as colored triangles. (**C**) Representative par^2^FISH images for several top variable genes, with sample OD_600_ values indicated below each panel. Fluorescence intensity is shown via the inferno color bar.

Several notable features were highly conserved across species. For example, the porins *ompC* and *ompD* also exhibited subpopulation-specific expression in *S. Typhimurium*, further supporting potentially regulated differences of outer membrane permeability among individuals sharing the same well-mixed environment (Fig. 4B–C). Both species also diversified cell-cell expression of maltose (*malE/lamB*) and sorbitol (*srlD*) utilization pathways, as in *E. coli*. However, *S. Typhimurium* further expanded this pattern to various other putative metabolites such as galacturonate (*uxuA*), glucarate (*garL*) 4-hydroxyphenylacetate (*hpaD*), and fructose (*fruK*) (Fig. 4B-C; Table S5). The absence of these potential substrates in our defined media and the emergence of these subpopulations at higher densities (Fig. 4B) hint at an evolutionary conserved bet-hedging strategy driven by the relaxation of catabolite repression.

In other cases, similar processes, rather than specific genes, showed hypervariable expression. For example, an evolutionarily distinct *fimA* gene (*40*), which is not under regulation by a known phase variation mechanism (*41*), was nonetheless the most hypervariable locus in *S. Typhimurium* (Fig. 4B). These data indicate that regulating subpopulation-level adhesion is a conserved strategy executed through potentially different molecular mechanisms. Various iron acquisition systems also displayed subpopulation-specific expression in each species, including ferric catecholate siderophore transporters (*cirA*, *fiu*) in *E. coli* and enterobactin biosynthesis (*entB*) and uptake (*fepA*) genes in *S. Typhimurium*. Comparable equivalencies were also observed for various genes involved in amino acid biosynthesis, nitrogen metabolism, and DNA damage responses (Tables S2 and S5).

In summary, this first-of-its-kind comparative analysis reveals that the phenotypic landscapes of *E. coli* and *S. Typhimurium* share a closely aligned functional structure, while highlighting processes and genes that exhibit either striking conservation or species-specific divergence.

### Specialization and environmental control of virulence heterogeneity under infection-related conditions

*S. Typhimurium* exhibits species-specific virulence programs that are deployed during epithelial invasion and intracellular growth. To quantify cell-to-cell heterogeneity in these programs under infection-relevant conditions, we extended par^2^FISH to conditions that induce the two major pathogenicity islands, SPI-1 in ESP in rich LB medium and SPI-2 in media mimicking the intracellular state (*42*, *43*). Imaging more than 160,000 cells from seven additional samples, we identified numerous hypervariable genes, many of which were also detected in our M9 experiment or highlighted in recent MATQ-seq analysis under similar conditions (*17*) (Table S6). Notably, subpopulations expressing diverse carbon utilization genes were also detected in LB, again peaking at early stationary phase when nutrients become limited (fig. S6A; Table S6). Considering global levels of heterogeneity, median EVI scores were highly similar across media and growth phases, indicating that overall heterogeneity is largely robust to environmental and physiological changes at the mRNA level (Fig. 5A).

**Fig. 5.**
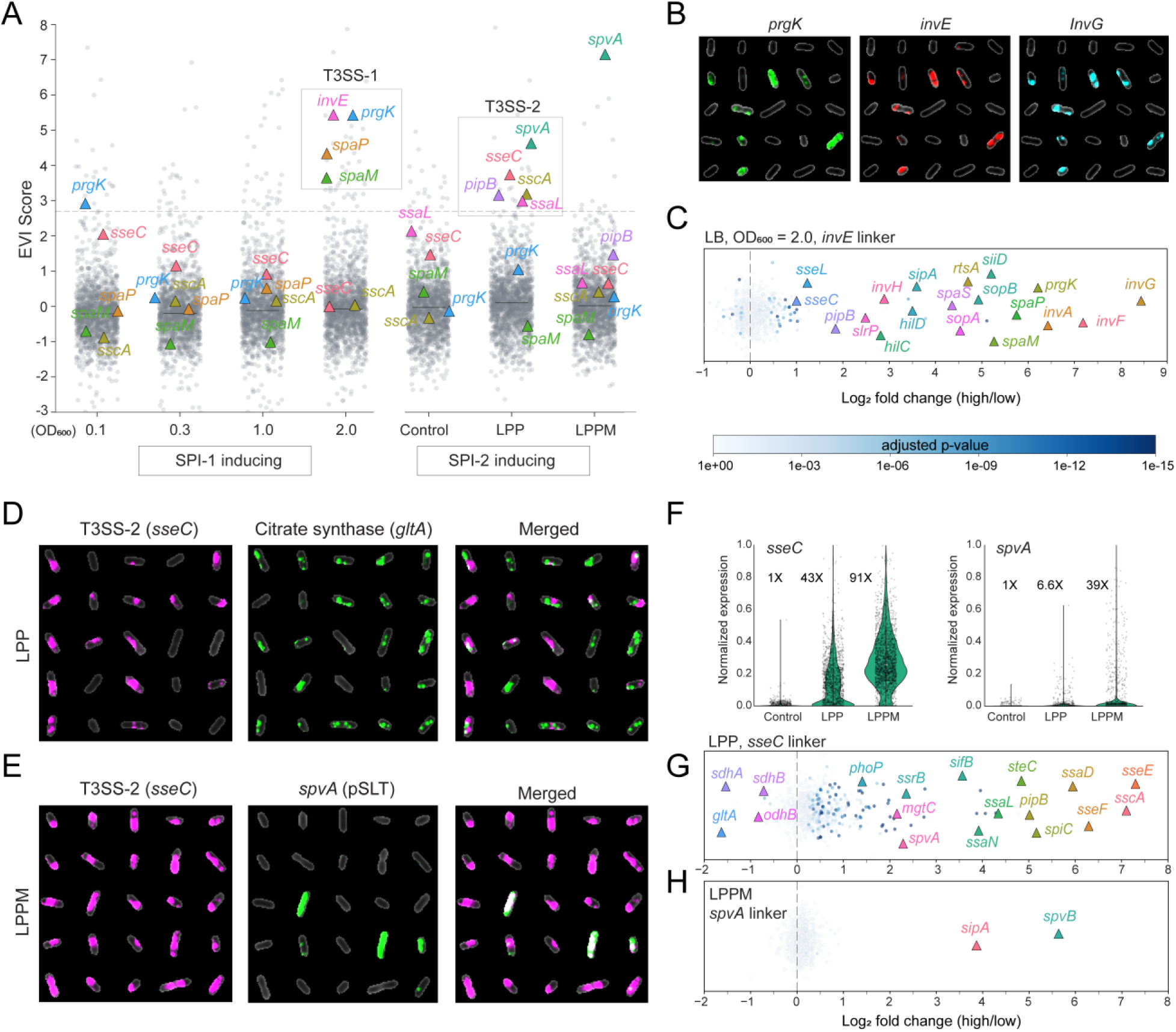
Control of virulence heterogeneity under infection-related conditions. (**A**) Distribution of EVI values under SPI-1– and SPI-2–inducing conditions, as indicated on the x-axis. The hypervariability threshold EVI (2.7) is shown as a gray dashed line. Median EVI per condition is shown as a horizontal black line; all gene–condition datapoints are shown in gray, with selected genes highlighted as colored triangles. (**B**) Representative par^2^FISH signals for several top-variable T3SS-1 genes in cells grown in LB to an OD_600_ of 2.0 (SPI-1–inducing). (**C**) FISH-Links analysis using the T3SS-1 marker gene *invE*. The x-axis shows the log_2_ fold-change in expression between *invE*^High^ and *invE*^Low^ subpopulations. Points are colored by adjusted *p*-value according to the color bar. Highly significant genes of interest are highlighted with colored triangles and labeled by name. (**D**) Cells grown in LPP medium overlaid with par^2^FISH signals for the T3SS-2 marker *sseC* (magenta) and the TCA cycle gene *gltA* (green). (**E**) Cells grown in LPPM medium overlaid with par^2^FISH signals for *sseC* (magenta) and *spvA* (green). (**F**) Violin plots of single-cell expression for *sseC* and *spvA*. Conditions are indicated on the x-axis. Expression was normalized to the maximum observed after removal of extreme outliers (top 99.9th percentile). Individual cell values are shown as black points. Average expression across all cells (normalized to the lowest value, 1X) is shown within each plot. (**G–H**) FISH-Links analysis using *sseC* and *spvA* in LPP and LPPM media, respectively, as indicated in each panel. Statistical significance of individual genes is color-coded according to the reference bar in panel C.

*S. Typhimurium* relies on the SPI-1 encoded type 3 secretion systems (T3SS-1) to invade intestinal epithelial cells and initiate infection (*44*). Consistent with previous reports, numerous T3SS-1 genes (e.g., *prgK*, *invA*, *spaP*) were expressed in a strongly bistable manner in LB cultures reaching ESP (OD_600_ = 2.0; Fig. 5A-B) (*45*). To characterize this subpopulation, we used the *invE* gene as a linker. We identified a T3SS-1^High^ subpopulation that was enriched 50-fold on average for diverse T3SS-1 structural genes, regulators (e.g., *invF*, *hilD*, etc.), and effectors (e.g., *sopB*), indicating full pathway activation within a minority subset (*44*) (Fig. 5C; Table S7). Adhesion to epithelial cells, mediated by SPI-4–encoded adhesins, has been shown to be co-regulated with SPI-1 via HilA co-regulation (*46*). In line with this, the SPI-4 marker *ssiD* was >33-fold enriched in the T3SS-1^High^ subpopulation (Fig. 5C; Table S7). Several genes of the SPI-2–encoded T3SS-2 (e.g., *sseL*, *sseC*, *pipB*) also showed induction in T3SS-1^High^ cells (up to ∼2-fold; Fig. 5B; Table S7). Indeed, such partial cross-activation has been previously reported, and depends on *hilD* (*47*). These results further validate the FISH-Links approach by capturing the full SPI-1 virulence program, including its known regulatory couplings, within a defined single-cell subpopulation. These results further validate the FISH-Links approach by resolving the canonical SPI-1 bistable state as a coherent, fully activated virulence program with its known regulatory couplings at single-cell resolution.

Following macrophage phagocytosis, *S. Typhimurium* deploys the SPI-2–encoded T3SS-2 to prevent phagosome–lysosome fusion and remodel the *Salmonella*-containing vacuole (SCV) (*48*). T3SS-2 expression is triggered primarily by SCV acidification and via phosphate and Mg^2+^ depletion, which provide synergistic signals that enhance and sustain induction (*49*). To dissect the contribution of these cues at the single-cell level, we used SPI-2–inducing conditions based on low pH and phosphate (LPP) and a corresponding medium with reduced magnesium (LPPM). Strikingly, T3SS-2 genes exhibited a strongly bistable expression pattern, specifically under LPP conditions, with approximately 60% of the population at an induced state (Fig. 5A, D). In contrast, nearly all cells expressed T3SS-2 in LPPM, resulting in a largely uniform activation (Fig. 5A, E). These condition-specific patterns of bistability and uniformity were independently confirmed using an SPI-2 promoter reporter (fig. S7). In absolute terms, T3SS-2 marker mRNAs (e.g., *sseC*, *sseE*, *sscA*) ranked among the most abundant transcripts, reaching levels comparable to some ribosomal protein genes (e.g., *rplV*), underscoring the magnitude of this induction. At the population level, average T3SS-2 expression increased >40-fold in LPP relative to the control and more than 90-fold in LPPM (Fig. 5F). Thus, the difference between LPP and LPPM largely reflects a shift in the fraction of T3SS-2 induced cells rather than in activated-cell expression levels. These data suggest *S. Typhimurium* modulates virulence activation rates to match environmental uncertainty, in agreement with a recent report (*50*).

FISH-Links analysis using the *sseC* T3SS-2 marker revealed extensive induction of the structural and effector repertoire of this system in induced cells (Fig. 5G). Key positive regulators such as *phoP* and *ssrB* were upregulated 2.7 and 5-fold, respectively (Table S7). Strikingly, genes of the tricarboxylic acid (TCA) cycle, including *gltA* (citrate synthase), *sdhA* (succinate dehydrogenase), and *odhB* (oxoglutarate dehydrogenase), were downregulated in induced cells (Fig. 5D and G). Previous studies showed that *gltA* and *sdhA* deletion mutants replicate more efficiently in resting and activated murine macrophages, and proteomic analyses of intracellular *S. Typhimurium* similarly revealed suppression of TCA cycle components (*51*, *52*). Together, these single-cell transcriptomic data indicate that T3SS-2 activation is coupled to coordinated remodeling of central metabolism within the same subpopulation, linking virulence induction to metabolic reprogramming associated with intracellular survival.

The pSLT virulence plasmid of *S. Typhimurium* encodes several regulators and effectors that complement chromosomal pathogenicity islands during infection (*53*, *54*). Among these, we quantified the expression of *spvA* and *spvB* from the *spvRABCD* operon, which encodes several effectors that contribute to intracellular survival via T3SS-2–associated functions (*53*). Whereas magnesium depletion led to largely uniform activation of SPI-2 genes, *spvA* and *spvB* expression both increased and became significantly more heterogeneous, indicating that their regulation can be decoupled from canonical SPI-2 control (Fig. 5A, F; fig. S8). Consistent with this, FISH-Links analysis using *spvA* as an anchor under LPPM conditions enriched primarily for *spvB* and, to a lesser extent, the T3SS-1 effector *sipA* (Fig. 5H; Table S7), suggesting a largely closed and autonomous regulatory module at the *spv* locus. In contrast, other effectors such as *pipB* closely tracked T3SS-2 structural gene expression, consistent with co-regulation (fig. S8; Table S7). These findings indicate that the *spvRABCD* operon is strongly expressed in only a subset of T3SS-activated cells, implying that secretion of these plasmid-encoded effectors is restricted to a minority subpopulation. This subpopulation-specific expression appears unique to the *spv* locus and may reflect plasmid-specific regulatory bistability or feedback control, highlighting diversification of virulence functions within an otherwise uniformly activated pathogenic program.

### Full-transcriptome profiling of T3SS-induced cells via FACS–RNA-seq

FISH-Links enables parallel profiling of multiple transcriptionally distinct subpopulations but is inherently limited to the predefined gene set targeted by par^2^FISH. In scRNA-seq workflows, unsupervised clustering groups cells with similar transcriptional states, enabling transcriptome-wide differential expression analysis between inferred subpopulations despite low mRNA capture efficiency per cell (*12–18*). Inspired by this conceptual framework, we sought to obtain a comprehensive transcriptomic characterization of defined bacterial subpopulations by physically isolating cells according to the expression of key marker genes. To this end, we optimized an approach combining marker-based FACS sorting with efficient bulk-RNA sequencing of as few as 10^7^ fixed cells. We constructed a dual-promoter reporter plasmid carrying an unstable P*sseL*::mChartreuse to label the T3SS-2^High^ subpopulation and P*rpsM*::mScarlet-I3 as a constitutive marker for all cells. Using this reporter, *S. Typhimurium* LPP cultures were sorted into T3SS-2^High^ and T3SS-2^Low^ subpopulations in three biological replicates (25 million cells per subpopulation) and subjected to RNA sequencing (Fig. 6A). Averaging the expression profiles of the sorted subpopulations accurately reconstructed the transcriptomic signature of an unsorted population control (*r* = 0.99; Fig. 6B), indicating that population unmixing yields reproducible, genome-wide expression profiles that reflect the transcriptional heterogeneity of the original population.

**Fig. 6.**
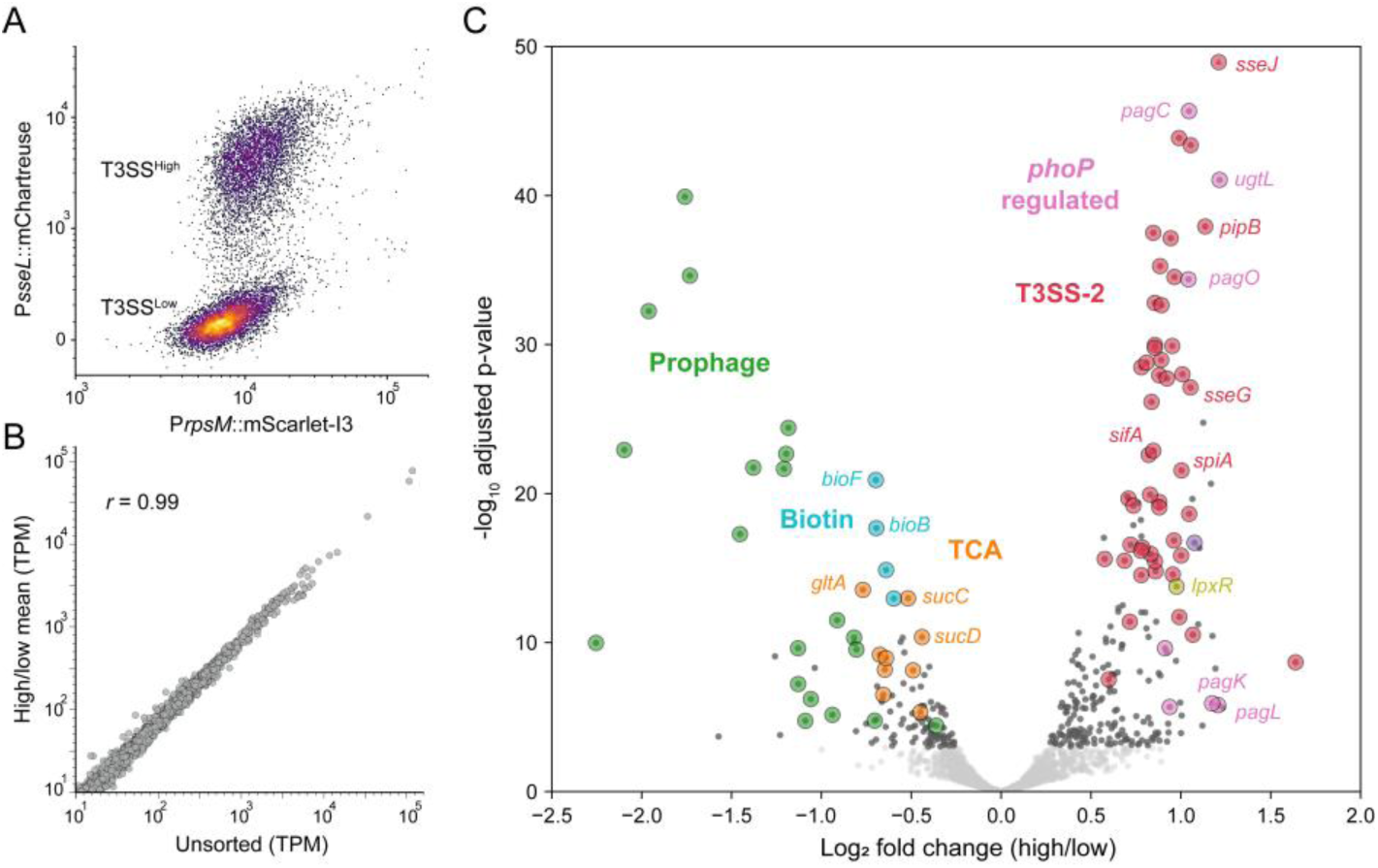
Deep transcriptomic profiling of FACS-sorted T3SS-2 subpopulations. (**A**) FACS scatter plot of *S. Typhimurium* cells harboring a dual-reporter plasmid, with unstable P*sseL*::mChartreuse-LVA reporting T3SS-2 expression and P*rpsM*::mScarlet-I3 labeling all cells. Axes are displayed using logicle transformation. T3SS-2–expressing (T3SS-high) and non-expressing (T3SS-low) subpopulations are indicated. (**B**) RNA-sequencing comparison of the unsorted population (x-axis) versus the combined mean expression of the T3SS-high and T3SS-low subpopulations (y-axis). Gene expression is shown in transcripts per million (TPM); genes with <10 TPM in all samples were excluded. The Pearson correlation coefficient is indicated in the figure. (**C**) Volcano plot showing differential gene expression between T3SS-high and T3SS-low cells for genes covered on average by at least 10 TPM. The x-axis indicates log_2_ fold change, and the y-axis shows the p-value (−log₁₀) adjusted for multiple hypothesis testing. Genes with adjusted p-values <0.001 are shown in light gray, while genes above this threshold are shown in dark gray. Selected genes of interest are further highlighted with circles colored according to functional pathways or processes, with gene names annotated in some cases (e.g., TCA genes shown in orange).

Differential expression analysis was highly concordant between the two approaches: significant genes were similarly up- or downregulated in both datasets (142/145; Fisher’s exact test, *p* < 10⁻^10^) and showed good correlation in fold-change magnitude (*r* = 0.72, *p* < 10⁻^10^). Importantly, FACS-RNA-seq indicated a global induction of SPI-2 genes and their key regulators (e.g., *ssrAB*, *phoP/Q*), together with repression of the TCA cycle genes detected above (e.g., *gltA*, *sdhA*, *odhB*) (Fig. 6C; Table S8). Differential expression of diverse metabolic genes was observed, consistent with metabolic reprogramming in T3SS-2 induced cells (Fig. 6A; Table S8). The T3SS-2^High^ subpopulation was also enriched for genes involved in lipopolysaccharide modification and immune evasion, including *lpxR* (also detected by FISH-Links; Tables S7–S8) and the PhoP-activated genes *pagL* and *pagO* (*55*) (Fig. 6C). This analysis also revealed genes not included in our par^2^FISH screen. For example, additional PhoP/Q-regulated virulence genes (*pagC*, *pagD*, *pagN*), as well as *ugtL*, which amplifies the PhoP/Q response (*56*). These data are consistent with differential PhoP/Q activity between subpopulations sharing the same well-mixed environment (Fig. 6C; Table S8). In contrast, the most strongly downregulated genes in T3SS-2^High^ cells mapped to two prophage regions, suggesting transcriptional silencing of foreign elements (Fig. 6C; Table S8). Several of these prophage genes were included in the par^2^FISH panel (and showed hypervariable expression in agreement with spontaneous induction; Tables S5 and S6) but were expressed in too few cells for FISH-Links analysis, further highlighting the complementary power of deep subpopulation RNA-sequencing.

## Discussion

Here, we introduce an integrated strategy for mapping bacterial phenotypic landscapes with high sensitivity and scale by combining the strengths of high-content imaging and RNA sequencing. Leveraging direct measures of expression variability, our approach highlights genes whose cell-to-cell variability far exceeds baseline expression noise levels, pointing to regulated states and phenotypic differentiation. Applying this framework to *E. coli* and *S. Typhimurium*, we uncover transcriptionally defined subpopulations with potential roles in metabolism, stress response, antibiotic tolerance, and virulence. These results complement existing single-cell datasets and can help guide downstream functional and mechanistic investigations.

By performing a comparative single-cell analysis under the same growth conditions and population densities, we show that for most genes, expression variability is well conserved between species and that extreme expression variability is a relatively rare property. When present, such hypervariability can be conserved either at the orthologue level or across different genes from functionally related pathways. Notably, certain behaviors, such as diversification of carbon utilization strategies, were observed in both species across multiple growth conditions. This conservation suggests that regulated metabolic heterogeneity may represent a broadly conserved bet-hedging strategy under environmental uncertainty, joining previous observations of complex resource utilization in isogenic bacterial populations (*10*, *35*, *36*). More generally, these data indicate that phenotypic heterogeneity is selectively associated with specific physiological functions. Extension of this comparative approach across species and contexts can help reveal which cellular processes bacteria preferentially deploy in subpopulations, a critical step toward understanding the organizing principles and evolutionary basis of phenotypic heterogeneity.

With the ability to map bacterial subpopulations and their full transcriptomes, an important next step is to directly link transcriptionally defined states to quantitative functional readouts, such as growth rate, metabolic output, or infection-specific functions. For example, integrating growth rate measurements with single-cell transcriptomes could help determine whether particular subpopulations arise in response to environmental stress or represent phenotypes associated with fitness trade-offs, providing insight into potential bet-hedging strategies. Beyond single-cell gene expression, par^2^FISH reports on cell morphology and can be readily combined with reporter proteins or chemical dyes. These capabilities provide a potential foundation for developing high-throughput platforms for functional characterization of bacterial subpopulations.

Finally, beyond mapping and phenotypic characterization, a key challenge will be to systematically identify the genetic and regulatory circuits that give rise to subpopulation structure. Developing integrated experimental and computational strategies to link observed cellular differentiation to underlying network architecture and specific regulatory elements will be essential for achieving a mechanistic understanding of how bacteria generate and control phenotypic heterogeneity.

## Methods

### Bacterial strains and growth conditions

*E. coli* strain MG1655 (K-12) and *S. Typhimurium* strain 14028S were grown aerobically at 37 °C with shaking at 250 rpm in M9 minimal medium (6 g/L Na_2_HPO_4_, 3 g/L KH_2_PO_4_, 0.5 g/L NaCl, 1 g/L NH_4_Cl, 0.1 mM CaCl_2_, 1 mM MgSO_4_, and 5 mg/L thiamine; pH 7.1–7.2). The medium was supplemented with 0.2% (w/v) glucose as the sole carbon source and a 1:100 dilution of a 100X trace elements solution (5 g/L EDTA, 498 mg/L FeCl_3_, 84 mg/L ZnCl_2_, 13 mg/L CuCl_2_·2H_2_O, 10 mg/L CoCl_2_·6H_2_O, 10 mg/L H_3_BO_3_, and 1.6 mg/L MnCl_2_·4H_2_O). For growth curve experiments, overnight cultures grown in M9 medium were washed once in fresh medium and diluted 1:100 into 100 mL fresh medium in 250 mL baffled flasks. Cultures were incubated at 37 °C with shaking at 250 rpm and sampled at defined optical densities (OD_600_), as indicated. Samples were immediately fixed in ice-cold 2% paraformaldehyde (PFA; Electron Microscopy Sciences, 15174) for 1.5 h on ice, washed twice with 1X phosphate-buffered saline (PBS; Invitrogen, AM9625), resuspended in 70% ethanol, and incubated at −20 °C for 24 h to permeabilize cells.

For SPI-1 inducing conditions, *S. Typhimurium* was grown overnight in 3 mL LB medium [10 g/L tryptone, 5 g/L yeast extract, 5 g/L NaCl] at 37°C with shaking at 220 rpm. The following morning, cultures were washed twice in 1X PBS and diluted 1:1,000 into 250 mL flasks containing 25 mL of the indicated medium. Cultures were incubated at 37°C with shaking at 220 rpm and collected at defined growth phases based on OD_600_, corresponding to varying levels of *Salmonella* Pathogenicity Island (SPI)-1 expression (*42*): early exponential phase (EEP; OD_600_ = 0.1), mid-exponential phase (MEP; OD_600_ = 0.3), late exponential phase (LEP; OD_600_ = 1.0), and early stationary phase (ESP; OD_600_ = 2.0). For SPI-2 inducing conditions (*42*, *43*): culturing was performed using phosphate carbon nitrogen (PCN) minimal medium [4 mM Tricine, 100 µM FeCl_3_, 376 µM K_2_SO_4_, 50 mM NaCl, 0.4% (w/v) glucose, 15 mM NH_4_Cl, 0.01 mM CaCl_2_, 10 nM Na_2_MoO_4_, 10 nM Na_2_SeO_3_, 4 nM H_3_BO_3_, 300 nM CoCl_2_, 100 nM CuSO_4_, 800 nM MnCl_2_, 1 nM ZnSO_4_, 80 mM MES (pH 5.8), 0.4 mM or 25 mM K_2_HPO_4_/KH_2_PO_4_, and 1 mM or 10 mM MgSO_4_] which mimics phagosomal conditions within macrophages, overnight cultures were prepared as described above. Three PCN variants were used: SPI2-non-inducing conditions [SPI2-control; pH 7.4, 25 mM phosphate, 1 mM MgSO_4_], SPI2-inducing [LPP; pH 5.8, 0.4 mM phosphate, 1 mM MgSO_4_], and LPP with lower magnesium [LPPM; pH 5.8, 0.4 mM phosphate, 10 µM MgSO_4_]. All cultures were harvested at OD_600_ = 0.3. Samples were immediately fixed in 1 mL 3.7% PFA for 30 min at room temperature with gentle agitation, followed by permeabilization in 70% ethanol overnight at 4°C.

### Probe design and synthesis

Primary probes were designed as previously described (*19*, *23*). Briefly, all 30-nucleotide (nt) sequences with GC content between 45% and 65% and lacking more than four consecutive identical bases were extracted from the target regions and aligned to the reference genome using BLAST. Sequences with nonspecific matches of ≥18 nt were discarded. Negative control probes were designed against the P1 phage genome (NC_005856.1) using the same criteria. For each gene analyzed, non-overlapping probes were randomly selected and flanked by overhangs containing two repeats of a secondary hybridization sequence, spaced by “AA”, complementary to a designated fluorescent readout probe (*23*). For each organism, a set of 16 probe libraries was designed to target either 150 or 120 genes, corresponding to *E. coli* (2,400 total genes; Table S1) and *S. Typhimurium* (1,920 total genes; Table S4), respectively. Genes were randomly selected but were limited in most cases to one selection per operon. Each library also included probes targeting a small set of shared quality control genes, as described above (linkers; Fig. 1A). Each target gene was covered by between 8 to 35 non-overlapping probes randomly selected from the gene-specific probe pool, corresponding to a range of 16 to 70 fluorophores per gene (average of about 40).

Primary probe libraries were synthesized as oligonucleotide array pools (Twist Bioscience). For *E. coli* probe libraries, 20-nucleotide flanking sequences were appended to each probe to enable amplification by polymerase chain reaction (PCR) (forward, 5′-TTTCGTCCGCGAGTGACCAG-3′; reverse, 5′-GCATCCCGACATGGACGTTG-3′). For *S. Typhimurium*, probe libraries were synthesized in pairs within each oligonucleotide pool and therefore carried distinct flanking sequences (forward-1, 5′-TTTCGTCCGCGAGTGACCAG-3′; reverse-1, 5′-GCATCCCGACATGGACGTTG-3′; forward-2, 5′- AAGCGCCACGAGTTGTCACG-3′; reverse-2, 5′-GCATCACTTTGGGCTCGGCT-3′). Each oligo pool was resuspended in 1× TE buffer (Integrated DNA Technologies, 11-05-01-05) to a final concentration of 1 ng/µL. A two-step PCR amplification was performed using the KAPA HiFi HotStart ReadyMix PCR Kit (Roche, KK2602). The first PCR used 1 µL of oligo pool as input, followed by a second PCR consisting of eight parallel reactions, each seeded with 1 µL of the first PCR product. Amplification was performed for 10 and 9 cycles, respectively, with an annealing temperature of 68 °C. Secondary PCR products were pooled, purified using the QIAquick PCR Purification Kit (Qiagen, 28104), and analyzed using the Agilent TapeStation (D1000 ScreenTape, 5067-5582). Three micrograms of purified PCR product were used as template for in vitro transcription (New England Biolabs, E2040S). RNA products were analyzed using the Agilent TapeStation (High Sensitivity RNA ScreenTape, 5067-5579), followed by reverse transcription (Thermo Fisher Scientific, EP0753). The resulting single-stranded DNA probes were treated with 1 M NaOH at 65 °C for 15 min to hydrolyze RNA templates, neutralized with 1 M acetic acid, and purified by ethanol precipitation.

### Coverslip functionalization

Coverslips were cleaned with 100% ethanol, air-dried, and coated by immersion in a 1% bind–silane solution (3-(trimethoxysilyl)propyl methacrylate; Sigma-Aldrich, M6514-25ML) prepared in 10% (v/v) acidic ethanol (pH 3.5) for 1 h at room temperature. Coverslips were washed three times with 100% ethanol and dried in an oven at 90 °C for 30 min. They were then incubated overnight at room temperature with 100 µg/mL poly-D-lysine (Sigma-Aldrich, P7280). The following morning, excess poly-D-lysine was removed by thorough rinsing with ultrapure water, and coverslips were dried with compressed air and used fresh for each experiment.

### Par^2^FISH

Cellular barcoding was performed using probes targeting distinct regions of the 16S rRNA (designed as described above). Barcode set 1 (BC1) was used to encode sample condition, and barcode set 2 (BC2) was used to encode probe library identity. In addition, a third universal 16S rRNA probe was included as a cellular reference. Each barcoding probe carried two distinct secondary hybridization sequences that together defined the barcode identity, except the universal probe as described below. Two replicates of each independently fixed sample (1.5 x 10^8^ cells per reaction) were collected into microcentrifuge tubes and pelleted by centrifugation (12,000 rpm, 1 min). Cell pellets were permeabilized in 200 µL lysozyme solution [0.5 µg/mL lysozyme (Sigma-Aldrich, L3790), 0.1 M Tris-HCl (pH 7.0), 0.05 M EDTA] for 15 min at 37 °C, followed by two washes with 100 µL 1X PBS. Each sample was then hybridized individually with a BC1 probe (replicates received identical probes) by resuspension in 20 µL nuclease-free water containing 30 nM of the designated 16S rRNA barcode probe, mixing with 30 µL prewarmed primary hybridization buffer [50% (v/v) formamide (Thermo Fisher, AM9342), 10% (w/v) dextran sulfate (Sigma-Aldrich, D8906), 3.34X SSC (Invitrogen, 15557)], and incubation at 37 °C for >16 h.

Samples were pelleted by addition of 100 µL 2X SSC and centrifugation (12,000 rpm, 1 min) to displace the viscous hybridization buffer, washed twice with 75 µL wash buffer [55% (v/v) formamide, 0.1% (v/v) Triton X-100 (Sigma-Aldrich, 93443) in 2X SSC], and incubated in 75 µL wash buffer for 30 min at room temperature to remove nonspecific binding. Samples were then washed twice with 100 µL 2X SSC and pooled in equal volumes into a new microcentrifuge tube. The pooled sample was divided into separate reactions, each hybridized with a specific gene probe library mixture, 12.5 nM of the appropriate BC2 barcode probe, and the universal 16S rRNA reference probes. As a negative control, one reaction was processed without a gene-targeting probe library. Each reaction was mixed with 30 µL prewarmed primary hybridization buffer and incubated at 37 °C for approximately 48 hours. Hybridized samples were washed as described above. Following hybridization, each sample was resuspended in 30 µL 4X SSC, and 10 µL from each reaction were combined to generate a final mixture. Ten microliters of this mixture were supplemented with 2 µL 0.01% Tween-20 to reduce cell clumping, and 4 µL were gently deposited at the center of a functionalized coverslip within a ring-shaped adhesive gasket. Samples were incubated at room temperature for 5 min to allow cell sedimentation and surface attachment. Coverslips were then centrifuged for 15 min at 500 rpm to form a uniform cell monolayer, followed by centrifugation at 4,000 rpm to stabilize surface binding.

Cells were immobilized in a hydrogel composed of 4% (v/v) acrylamide/bis-acrylamide solution (19:1; Bio-Rad, 1610154), 0.06 M Tris-HCl, and 0.5 M NaCl. The solution was degassed on ice under N_2_ for 15 min, after which polymerization was initiated by addition of 1 µL 250 mg/mL ammonium persulfate (APS; Sigma-Aldrich, A3678) and 1 µL N,N,N′,N′-tetramethylethylenediamine (TEMED; Bio-Rad, 1610800). Fifteen microliters of the reactive mixture were cast over the cell monolayer, covered with a clean glass slide, and degassed in a sealed container under N_2_ for 15 min. The container was then maintained at room temperature for an additional 20 min to allow complete gelation. Following polymerization, the slide and gasket were gently removed, and a flow chamber (Ibidi, 80168) was attached directly to the coverslip.

### Sequential imaging

Imaging was performed using a combined widefield microscopy and automated fluidics delivery system, as previously described (*23*). Image acquisition was carried out on a widefield inverted ECLIPSE Ti2-E microscope (Nikon) equipped with a motorized stage and controlled using the NIS-Elements software suite. Focus was maintained throughout imaging using the Perfect Focus System (PFS), and images were captured with a Prime BSI Express sCMOS camera (Teledyne). All par^2^FISH imaging was performed using a CFI Plan Apochromat 60X oil-immersion objective (NA 1.4; Nikon, MRD31605). Fluorophore excitation was provided by a SOLA III FISH Light Engine (Lumencor) in combination with standard fluorescence excitation/emission filter cubes for DAPI, FITC (Alexa Fluor 488), TRITC (Atto 550), and Cy5 (Atto 647). Imaging automation was implemented using the NIS-Elements JOBS module. Fluidic flow was driven by a CETONI Nemesys S module pump equipped with a 2.5 mL glass syringe. Reagent delivery was controlled via a Qmix valve module and a rotAXYS 360° positioning system (CETONI). Fluidics automation was operated using CETONI Elements software, which was coordinated with the NIS-Elements JOBS module to synchronize fluid exchange and image acquisition.

Chambered coverslips containing gel-embedded samples were mounted on the microscope and connected to the CETONI fluidics system. Regions of interest were identified by phase-contrast imaging. Sequential cycles of secondary probe hybridization, washing, and imaging were performed to acquire barcoding and mRNA fluorescence signals. Each hybridization round used a mixture of three distinct 15-nt readout probes, each labeled with Alexa Fluor 488 (A488; Integrated DNA Technologies), Atto 550 (A550; biomers.net), or Atto 647 (A647; biomers.net). Readout probe solutions were prepared in EC buffer [10% (w/v) ethylene carbonate (Sigma-Aldrich, E26258), 16.7% (w/v) dextran sulfate (Sigma-Aldrich, D4911), 3.33X SSC] containing 50 nM of each probe. Hybridization solutions were flowed into the imaging chamber and incubated for 25 min at room temperature to allow probe binding. Following hybridization, samples were washed with 10% formamide wash buffer [10% (v/v) formamide, 0.1% (v/v) Triton X-100, and 2X SSC] to remove excess and nonspecifically bound probes (two washes followed by a 5-min incubation), followed by washing in 2X SSC. DNA was stained with 4′,6-diamidino-2-phenylindole (DAPI) [10 µg/mL (Sigma-Aldrich, MBD0015) in 2X SSC]. Imaging was performed in an oxygen-scavenging, anti-bleaching buffer consisting of 50 mM Tris-HCl (pH 8), 300 mM NaCl, 0.525 mg/mL Trolox (Sigma-Aldrich, 238813), 0.8% (v/v) glucose (Sigma-Aldrich, 49163), catalase (1:1000 dilution; Sigma-Aldrich, C3155), 0.5 mg/mL glucose oxidase (Sigma-Aldrich, G2133), and 2X SSC.

After each imaging round, readout probes were stripped by incubation in 55% formamide wash buffer for 3 min, followed by rinsing with 2X SSC. Stripping was performed twice to ensure complete signal removal. The cycle of hybridization, imaging, and probe stripping was repeated over multiple rounds to record barcode identities, mRNA expression, and background fluorescence.

### Demultiplexing and gene expression measurement

Sequential images were registered using either phase-contrast or DAPI fluorescence to correct for mechanical drift between imaging rounds. Channel misalignment between fluorophores was corrected by image alignment using the universal 16S rRNA signal, which was detected in all three fluorescence channels (A488, A550, and A647). Cell segmentation was performed on phase-contrast images using Cellpose (v2.0.21). A quality-control filtering step was applied to remove segmentation artifacts and auto-fluorescent objects. Secondary probe specificity was evaluated using the negative control sample, and a small number of readout probes exhibiting nonspecific binding were excluded from downstream analysis.

For sample demultiplexing, 16S rRNA fluorescence intensities were background-subtracted and quantified on a per-cell basis for each barcode readout, and barcode identities were assigned by thresholding these signals. The signal from the universal 16S rRNA reference probe was used to normalize for intrinsic variability in baseline 16S rRNA abundance across conditions. The false classification rate was estimated by quantifying the fraction of cells assigned to barcode combinations not used in the experiment.

To quantify gene expression, mRNA-FISH spots were identified as regional intensity maxima following application of top-hat filtering. Spots were then filtered by intensity thresholds defined using negative control genes and guided by the no-library control to ensure high specificity. The integrated intensity of each spot was calculated by summing pixel values within a 3×3 window centered on the local maximum. Detected spots were assigned to individual cells based on segmentation masks, and the total fluorescence signal was summed per gene per cell. Total mRNA-FISH fluorescence intensity was normalized by the number of probes designed for each gene.

### Expression variability analysis

Mean expression, coefficient of variation (CV), and tails expression ratio (TER; defined as the ratio between the mean expression of the top and bottom 5% of expressing cells) were computed for each gene under each condition in each experiment. Genes detected in fewer than 75 cells or exhibiting extreme outlier standard deviations were excluded from further analysis. Remaining genes were grouped by fluorophore (A488, A550, and A647), and the relationships between log-transformed mean expression and log-transformed CV or TER were modeled independently within each group using third-degree polynomial regression. Model parameters were estimated by ordinary least squares using the statsmodels Python package. A 95th-percentile confidence interval was derived from each fitted relationship, and residuals for CV (CVR) and TER (TERR) were calculated as the difference between each observed value and its corresponding predicted interval-bound value. Residuals were then standardized (z-scored) per metric and averaged to generate a unified expression variability index (EVI). Hypervariable genes were defined as gene–condition pairs with z-scores exceeding 1.96 for both CVR and TERR. Final EVI thresholds were applied as indicated above (*E. coli* M9, EVI = 2.7; *S.* Typhimurium M9, EVI = 2.21; *S. Typhimurium* SPI-inducing, EVI = 2.7). For the comparative analysis between E. coli and S. Typhimurium, a set of 826 orthologous genes was identified with EcoCyc (*57*). The maximal EVI score for each gene was compared as described.

### FISH-Links

For the *E. coli* M9 experiment, linker gene probes were included in the original set of 16 probe libraries (Table S3). For the *S. Typhimurium* experiments under SPI-inducing conditions, a second stage par^2^FISH experiment was performed using newly designed probes targeting *invE*, *sseC*, *pipB*, and *spvA* (Table S7). These probes were synthesized as a ready-to-use oligonucleotide pool (Integrated DNA Technologies) and added to each reaction as exogenous linker probes.

To derive transcriptional signatures of subpopulations, differential gene expression analysis was performed by stratifying cells based on linker gene expression measured in each cell. For each sample, cells in the top 5% and bottom 5% of linker expression were classified as Linker^High^ and Linker^Low^ subpopulations, respectively. In practice, due to the zero-inflated nature of bacterial single-cell mRNA measurements, the Linker^Low^ subpopulation was sometimes defined as all cells with no detectable expression of the linker gene. Average gene expression was quantified for genes detected in at least 30 cells per group and with minimal expression exceeding 20 fluorescence units in at least one group. Expression values were normalized by cell volume, calculated by approximating cells as spherocylinders with dimensions defined by the major (long) and minor (short) axes. For each gene, log₂ fold changes in expression between Linker^High^ and Linker^Low^ subpopulations were computed, and statistical significance was assessed using a two-sided Mann–Whitney U test. P-values were corrected for multiple hypothesis testing using the Benjamini–Hochberg procedure with a false discovery rate (FDR) of 0.01.

### Promoter reporter imaging

Publicly available promoter reporter strains (*58*) were used to validate par^2^FISH-identified hypervariable genes in *E. coli*. Reporter strains were grown in M9 medium supplemented with 0.2% glucose, as described above, with the addition of kanamycin (25 µg/mL), and imaged on 1% agarose pads upon reaching the OD_600_ values indicated in the figures (Fig. 2E; Fig. S3). Images were acquired using a CFI Plan Apochromat 100X oil-immersion objective (NA 1.45; Nikon, MRD01905). Fluorescence excitation was provided by either a SOLA III FISH Light Engine (Lumencor) or a pE-800 LED illumination system (CoolLED, pE-800-L-MB-SYS-ZZ).

For *S. Typhimurium*, a *PsseL*::CFP promoter reporter was constructed by Gibson assembly using pSEVA237C (GenBank accession JX560403) as the backbone vector. The backbone plasmid was linearized by digestion with NdeI-HF (New England Biolabs, R3131S). The *sseL* promoter fragment was amplified by colony PCR from *S. Typhimurium* genomic DNA using primers sseL_FW (5′-GCAGGCATGCAAGCTTAGGAGGAAAAACATCTCTTGTATCGACGCGTTACCAG-3′) and sseL_REV (5′- GCTCCTCGCCCTTGCTCACCATGTTCTCTACGGCGCTAAAC-3′). Correct assembly was confirmed by Sanger sequencing.

### FACS-RNA-seq

For FACS-RNA-seq experiments, we constructed plasmid p15A_pSseL-mChartreuse-LVA_pRpsM-mScarlet-I3. This dual-fluorescence reporter plasmid contains the *sseL* promoter driving expression of destabilized mChartreuse (*59*) as a SPI-2 activity reporter, and the constitutive *rpsM* promoter driving mScarlet-I3 (*60*) expression to label all cells. The mChartreuse protein was destabilized by addition of a C-terminal LVA degradation tag (*61*) to improve the temporal resolution of promoter activity measurements. The plasmid was synthesized commercially (Twist Bioscience).

S. *Typhimurium* carrying the P*sseL*::mChartreuse/P*rpsM*::mScarlet-I3 reporter were grown in LPP media as described above, with the addition of 100 µg/mL ampicillin. Cells were then fixed in PFA 3.7% and stored in 70% EtOH. Sorting was carried out on a BD FACSAria III cell sorter fitted with a 70 µm nozzle, a Neutral Density Filter of 1, and operated in four-way purity mode. mChartreuse signal detection was achieved using 488 nm laser excitation with emission collected through a 530/30 nm filter. mScarlet signal detection was achieved using 561 nm laser excitation with emission collected through a 582/15 nm filter. Events were acquired using BD FACSDiva software (v8.0), with forward scatter (FSC) and side scatter (SSC) thresholds set to 700 to limit background signal. The primary cell population was identified based on FSC and SSC characteristics, and as mScarelt positive, after which cells were partitioned into mChartreuse-high and mChartreuse-low subpopulations according to mChartreuse fluorescence intensity. Sorting was conducted at a flow rate of 1, yielding an event rate of approximately 9,000 events per second. For bulk samples, ∼5 x 10⁶ cells were deposited into 15 mL conical tubes containing 700 µL of 1X PBS, resulting in a final volume of approximately 6 mL. Flow cytometry data were processed and analyzed using FlowJo software and with the FlowCal python package.

For RNA extraction, approximately 2.5 x 10^7^ sorted cells (6 mL per 5 x 10^6^ cells) were centrifuged at 12,000 rpm for 20 min at 4 °C. Following centrifugation, the supernatant volume was reduced to ∼500 µL, and the pellet was resuspended by vortexing for 5 min prior to transfer into a 1.5 mL microcentrifuge tube. Samples were centrifuged a second time at 12,000 rpm for 7 min, after which the remaining supernatant was removed, leaving ∼50 µL. Cells were lysed by resuspension in 50 µL of lysis buffer composed of 7 M urea (Sigma-Aldrich, 57136), 0.1 M Tris-HCl pH 8.0 (Invitrogen, 15568025), 0.01 M EDTA disodium salt (Sigma-Merck, 03690), and 0.05 M NaCl (Invitrogen, AM9759), supplemented with 1 mg/mL lysozyme (Sigma-Aldrich, L6876). Samples were gently mixed and incubated at room temperature for 5 min. An additional 50 µL of the same buffer containing 1.2 mg/mL proteinase K (Rhenium, 25530049) was added, followed by incubation at 37 °C for 30 min. After enzymatic digestion, 650 µL of TRIzol reagent (Sigma-Merck, MFCD00213058) was added, and total RNA was purified using the Direct-zol RNA Miniprep Plus Kit (Zymo Research, R2070) in accordance with the manufacturer’s guidelines and after pooling the reactions. RNA concentration and integrity were evaluated using the Qubit RNA High Sensitivity Assay (Invitrogen, Q32855) and High Sensitivity RNA ScreenTape (Agilent, 5067-55979).

Ribosomal RNA depletion was performed using the Illumina Ribo-Zero Plus rRNA Depletion Kit (Illumina, 20040526) according to the manufacturer’s recommendations. Depleted RNA was used for library construction with the NEBNext® Ultra™ II Directional RNA Library Prep Kit for Illumina (NEB, E7760S), following the protocol for rRNA-depleted, degraded, or FFPE RNA samples. Library concentrations were measured using the Qubit dsDNA High Sensitivity Assay (Invitrogen, Q32855), and fragment size distributions were assessed using D1000 ScreenTape (Agilent, 5067-5590).

Sequencing was performed on an Illumina NovaSeq platform using 150 bp paired-end reads. RNA-seq data (read 1) were quality filtered and trimmed with fastp (*62*), aligned to the *S. Typhimurium* reference genome using Bowtie2 (*63*), and filtered by mapping quality. Duplicate reads were removed, and gene-level counts were generated using Subread (*64*) with strand-specific settings. Multimapping reads were discarded. Differential expression was performed using DEseq2 (*65*).

## Acknowledgments

We thank Danielle Lange, Roy Jacobson, and Irit Sherman, for critically reading and commenting on the manuscript, and members of the Dar laboratory for fruitful discussions and support. D.D. was generously supported by the ERC-StG programme (grant no. 101117863), the Israel Science Foundation (personal grant 2227/22), and the Alon fellowship. R.A. was generously supported by the Israel Science Foundation (grant no. 1289/22), Minerva foundation with funding from the Federal German Ministry for Education and Research, the Joint DFG-ISF Research Grant Program (grant no. 1297/25), the Pasteur–Weizmann Challenge, Knell Family Center for Microbiology, and the Shimon and Golde Picker - Weizmann Annual Grant. C.C.M. received funding from the European Union’s Horizon 2020 research and innovation programme under the Marie Sködowska-Curie grant no. 898715.

## Author contributions

Conceptualization: Z.P., D.D.

Methodology: Z.P., C.C.M., K.Z.G., O.A., D.D.

Investigation: Z.P., C.C.M., K.Z.G., R.A., D.D.

Formal analysis: Z.P., D.D.

Funding acquisition: D.D.

Supervision: D.D.

Writing – original draft: Z.P., C.C.M., K.Z.G, R.A., D.D.

## Data and materials availability

All data required for reproducing the figures and conclusions of this manuscript are presented in the main text or the supplementary materials. Raw data and scripts will be deposited.

