## Supplementary material for "Single-cell transcriptome imaging reveals conserved and virulence-linked phenotypic states in bacteria": SI

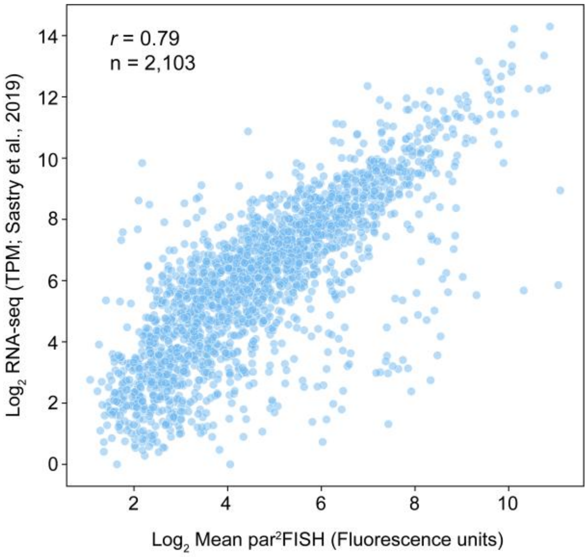


**fig. S1. Correlation of Par²FISH and bulk RNA-seq.** Scatter plot comparing par²FISH-derived mean expression levels ($OD_{600}$ = 0.75) with previously published bulk RNA-seq data (*1*). Shown are $log_2$-transformed mean fluorescence intensities (arbitrary units) versus normalized $log_2$-transformed RNA-seq counts. The number of analyzed genes and Pearson correlation value are shown within the figure.

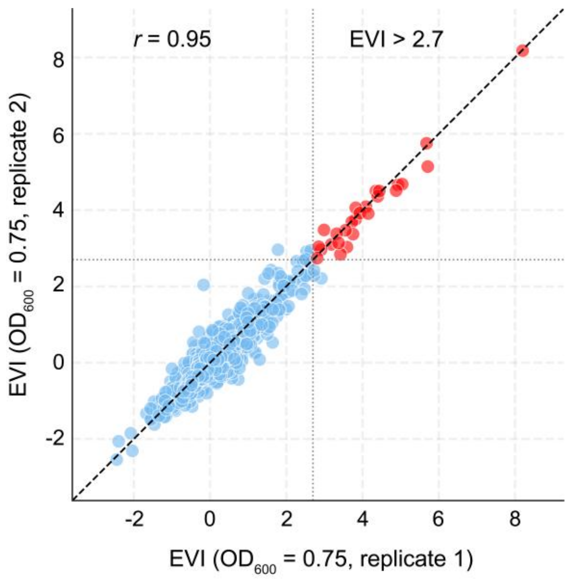


**fig. S2. Expression Variability Index (EVI) scores are highly reproducible across biological replicates.** Scatter plot comparing EVI scores of genes measured at $OD_{600}$ = 0.75 in two biological replicates. The Pearson correlation value is shown in the panel. The dashed lines indicate the EVI threshold for hypervariability (EVI = 2.7). Genes above and below this cutoff are highlighted in red and blue, respectively.

| Functional category | Hypervariable genes | Previously described |
|---|---|---|
| Acetate utilization | *aceB* | |
| Acid Resistance | *gadA, gadB, gadC, hdeD* | Urchueguía A., et al (2021),<br>McNulty, R., et al (2023) |
| Acid Resistance / carbon starvation | *slp* | Silander OK., et al (2012),<br>Urchueguía A., et al (2021) |
| Adhesion / Biofilm Formation | *flu (Ag43)* | Diderichsen B. (1980),<br>Henderson, I. R., et al (1997) |
| Arginine/pyrimidine biosynthesis | *carA, carB* | McNulty, R., et al (2023) |
| Arginine biosynthesis | *argA, argB, argC, argF, argG, argH* | McNulty, R., et al (2023) |
| Arginine transport | *artJ* | McNulty, R., et al (2023) |
| Aromatic amino acid biosynthesis | *aroF* | |
| Biotin biosynthesis | *bioB* | |
| Cell adhesion | *fimA* | Abraham J.M., et al (1985),<br>McNulty, R., et al (2023) |
| Colanic acid biosynthesis | *gmd, wcaI* | |
| Deoxyribonucleotide biosynthesis | *nrdA* | |
| Flagellar motility | *fliC* | McNulty, R., et al (2023) |
| Iron Acquisition | *cirA, fiu* | |
| Maltose utilization | *lamB, malE* | Sarfatis A., et al (2025) |
| Methionine biosynthesis | *metE, metF* | |
| Outer membrane porin | *ompC* | |
| Putrescine utilization | *puuA, puuB* | Sarfatis A., et al (2025) |
| Pyrimidine biosynthesis | *pyrB, pyrI* | Urchueguía A., et al (2021),<br>McNulty, R., et al (2023) |
| Ribosomal subunit protein | *rpsC* | |
| Sorbitol utilization | *srlD* | Urchueguía A., et al (2021) |
| SOS response | *recN* | Silander OK., et al (2012) |
| Sulfur assimilation | *cysJ, cysD* | Sarfatis A., et al (2025) |
| Uracil transport | *uraA* | |
| Zinc ion transport | *znuA* | |

**Table S2. Hypervariable genes detected in *E. coli*.** Genes with Expression Variability Index (EVI) scores above the hypervariability threshold in at least one M9 + 0.2% glucose sample. Genes are grouped by broad functional categories. Supporting literature demonstrating similar heterogeneity is cited where available.

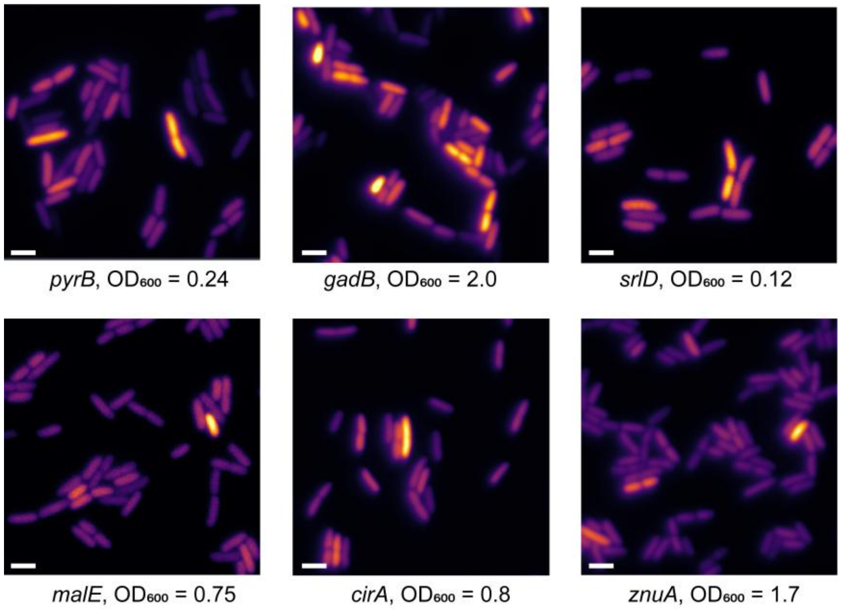


**fig. S3. Validation of hypervariable expression using promoter reporter strains.** Representative fluorescence microscopy images of six GFP promoter reporter strains grown in M9 + 0.2% glucose. The corresponding culture $OD_{600}$ values are indicated in each image. Scale bar, 2 μm

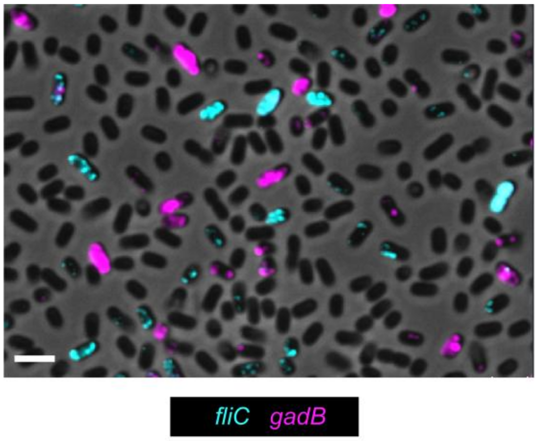


**fig. S4. Mutually exclusive expression of *fliC* and *gadB* in early stationary-phase LB cultures.** *E. coli* cells were grown in LB medium to $OD_{600}$ = 3.5 and imaged by phase-contrast microscopy. Overlaid are non-multiplexed smFISH fluorescence data for probes targeting *fliC* (cyan) and *gadB* (magenta). Scale bar, 2 µm.

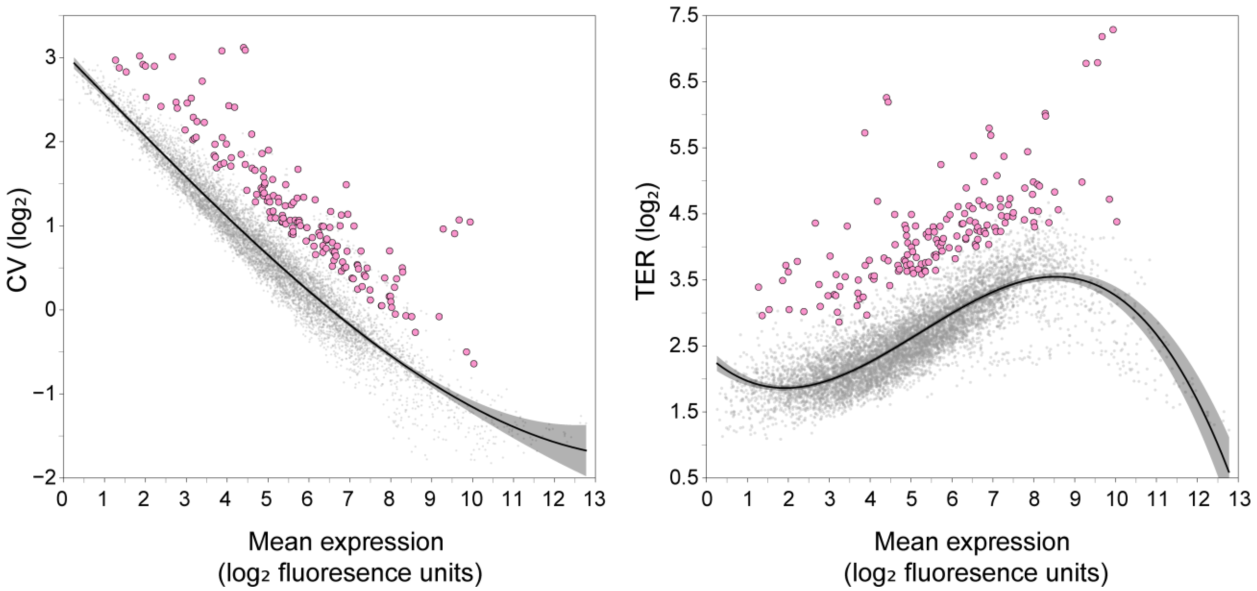


**fig. S5. Variability analysis in *S. Typhimurium*.** Coefficient of variation (CV) and Tails Expression Ratio (TER) values plotted against mean expression for each gene–condition datapoint. Individual datapoints are shown in gray, and those passing the heterogeneity threshold based on integrated CV and TER metrics are highlighted in pink. Mean expression values are reported in arbitrary fluorescence units. The fitted mean trend and 95% confidence intervals are shown as a black line and gray shading, respectively.

| Functional category | Hypervariable genes |
|---|---|
| 4-HPA catabolism | *hpaE, hpaD* |
| Aldolase | *STM14_RS15620* |
| Ammonium transport | *amtB* |
| Anaerobic respiration | *narJ, narI, nirC, nirB* |
| Aromatic amino acid biosynthesis | *aroF* |
| Ascorbate transporter | *STM14_RS12930* |
| Biotin biosynthesis | *bioD (STM14_RS04610)* |
| Cell adhesion | *fimA* |
| Chemotaxis | *cheA, cheB, cheW* |
| Enterobactin biosynthesis | *entB* |
| Enterohemolysin | *STM14_RS05495* |
| Flagellar motility | *flgB, flgD, flgF, flgJ, flgK, flgL, flgN, fliC, fliD, fliI, fliK, fliN ,fliP, fliS, fljB, motA, motB* |
| Fructose catabolism | *fruA, fruB, fruK* |
| GABA catabolism | *gabT* |
| Glucarate catabolism | *garL* |
| Glucosaminate catabolism | *dgaE* |
| Glutamine transport | *glnQ* |
| Hexuronate catabolism | *uxuA* |
| Hypothetical protein | *STM14_RS10975* |
| LPS modification | *pmrG* |
| Maltose utilization | *lamB, malM, malE* |
| Methionine biosynthesis | *metF* |
| Osmotic stress response | *osmY* |
| Outer membrane porin | *ompC, ompD* |
| Plasmid encoded gene (pSLT) | *STM14_RS00255* |
| Prophage | *STM14_RS11110, STM14_RS11005, STM14_RS10970, STM14_RS10945* |
| Purine biosynthesis | *STM14_RS13420* |
| Pyridoxal phosphatase | *cof* |
| Pyridoxal phosphate-dependent enzyme | *STM14_RS23305* |
| Quorum sensing | *lsrR* |
| Ribosomal subunit protein | *rpsC* |
| RNA helicase / Cold shock response | *deaD* |
| Siderophore receptor | *fepA* |
| SOS response | *sulA, umuC, sbmC* |
| SPI-1 virulence | *invE, sipC, sopB* |
| SPI-2 virulence | *sscA, spvA, pipB2* |
| Sulfur assimilation | *cysH* |
| T6SS | *STM14_RS14920* |
| Temperature-sensing protein (pSLT) | *tlpA* |
| Tricarboxylate transport | *STM14_RS14955* |

**Table S5. Hypervariable genes detected in *S. Typhimurium* grown in minimal media.** Genes with Expression Variability Index (EVI) scores above the hypervariability threshold in at least one M9 + 0.2% glucose sample. Genes are grouped by broad functional categories.

| Category | Hypervariable genes |
|---|---|
| Ammonium transport | *amtB* |
| Respiration / fermentation | *nirC, hyaD, hycF, fdnI,cydB2, narJ* |
| Amino acid biosynthesis and transport | *aroF, lysA, STM14_RS12100* |
| General stress | *dps, uspC* |
| Biotin biosynthesis | *bioD2* |
| Cell adhesion | *fimA* |
| Chemotaxis | *cheA* |
| Flagellar motility | *flgB, flgD, flgF, flgK, fliC, fliD, fliK, fliN, fljB, motB* |
| Fructose catabolism | *fruB, fruK* |
| Glucarate catabolism | *garL* |
| Gluconate catabolism | STM14_RS15605, idnK |
| Sorbitol catabolism | *srlD* |
| Hexuronate catabolism | *uxuA, uxaC* |
| Sialic acid catabolism | *nanA* |
| Lactate transport | *lldP* |
| LPS modification | *STM14_RS07500 (lpxR)* |
| Maltose utilization | *lamB, malM, malE, malP* |
| Osmotic stress | *osmY, osmE* |
| Outer membrane porin | *ompC, ompD, STM14_RS08505 (phoE)* |
| Prophage | *STM14_RS10940, STM14_RS11010, STM14_RS11020, STM14_RS11090, STM14_RS05730, STM14_RS05495* |
| Siderophores / iron acquisition | *STM14_RS03530 (FepA), iroB, entB* |
| Purine biosynthesis | *cof* |
| 1,2-propanediol catabolism | *pduL* |
| SOS response | *sulA, sbmC, recN* |
| SPI-1 virulence | *invE, prgK, spaM, spaP, spaR, invA, hilC* |
| SPI-2 virulence | *sscA, spvA, spvB, pipB, ssaL, sseC, STM14_RS07905* |
| Galactarate catabolism | *STM14_RS19555* |
| Temperature sensing protein (pSLT) | *tlpA* |
| Acid resistance | *cadA, cadB* |
| Misc. | *denD, fraD, htpG, phoU, STM14_RS00330, STM14_RS02415, STM14_RS03395, STM14_RS04105, STM14_RS06135, STM14_RS06785, STM14_RS07145, STM14_RS08275, STM14_RS10890, STM14_RS15620, STM14_RS11160, STM14_RS15580, STM14_RS16925, STM14_RS19870* |

**Table S6. Hypervariable genes detected in *S. Typhimurium* under SPI-inducing conditions.** Genes with Expression Variability Index (EVI) scores above the hypervariability threshold in at least one condition. Genes are grouped by broad functional categories.

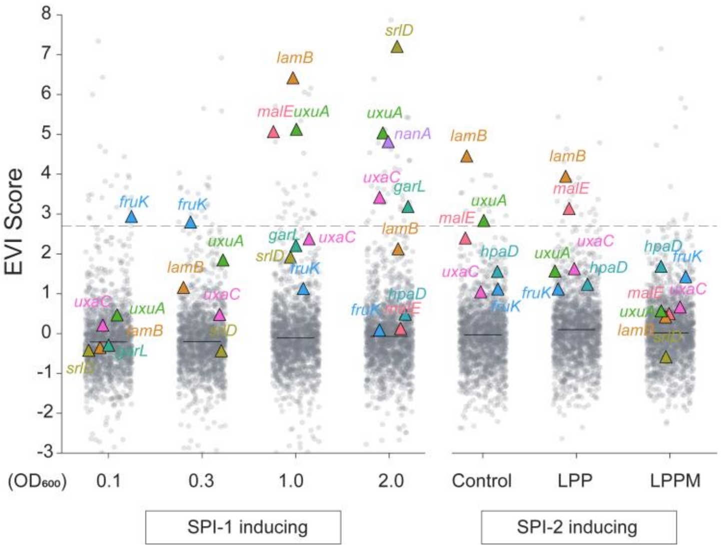


**fig. S6. Hypervariability of carbon utilization genes detected under virulence-inducing conditions.** Distribution of EVI values per condition (EVI calculated using datapoints from all conditions). The dashed line marks the EVI threshold (EVI = 2.7). Median EVI per condition is shown as a horizontal black line; all gene–condition datapoints are shown in gray, with various carbon source catabolism genes highlighted as colored triangles.

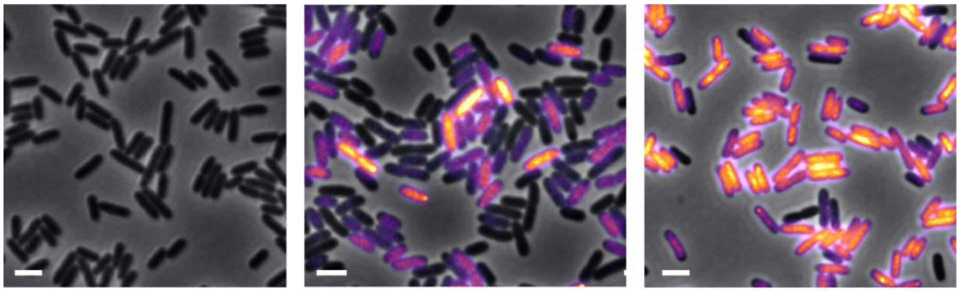


**fig. S7. Reporter strain validation of T3SS-2 expression heterogeneity.** *S. Typhimurium* T3SS-2 promoter reporter strain (P*sseL*::*cfp*) was grown under control, LPP, or LPPM conditions to $OD_{600}$ = 0.3 (as analyzed via par$^2$FISH) and imaged by fluorescence microscopy. Phase-contrast images are overlaid with CFP fluorescence (inferno color map), with intensity equalized across conditions for direct comparison. Scale bar, 2 μm.

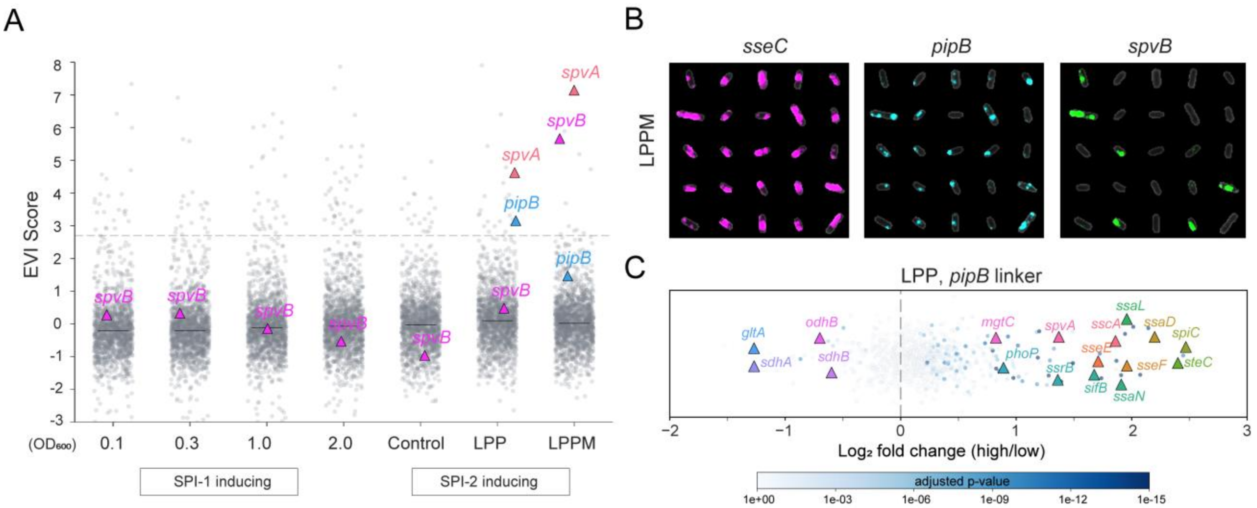


**fig. S8. T3SS-2 effectors display different heterogeneity regimes.** (**A**) Distribution of EVI values per condition (EVI calculated using datapoints from all conditions). The dashed line marks the EVI threshold (EVI = 2.7). Median EVI per condition is shown as a horizontal black line; all gene–condition datapoints are shown in gray, with effectors encoded on the *S. Typhimurium* chromosome (*pipB*) or pSLT virulence plasmid (*spvA* and *spvB*) highlighted as colored triangles. (**B**) Cells grown in LPP medium overlaid with par$^2$FISH signals for the T3SS-2 marker *sseC* (magenta), *pipB* (cyan) or *spvB* (green). (**C**) FISH-Links analysis using the effector gene, *pipB*. The x-axis shows the $\log_2$ fold-change in expression between *pipB*$^{High}$ and *pipB*$^{Low}$ subpopulations. Points are colored by adjusted *p*-value according to the color bar. Highly significant genes of interest are highlighted with colored triangles and labeled by name.
